# Time-restricted feeding near dawn entrains long-term behavioral changes through the suprachiasmatic nucleus

**DOI:** 10.1101/2021.02.18.431900

**Authors:** Qiaocheng Zhai, Yizhun Zeng, Yue Gu, Tao Zhang, Baoshi Yuan, Tao Wang, Jie Yan, Han Qin, Ling Yang, Xiaowei Chen, Antonio Vidal-Puig, Ying Xu

## Abstract

The suprachiasmatic nucleus (SCN) is a master circadian pacemaker known to integrate light intensity and seasonal information with peripheral tissues to coordinate daily rhythms of physiology and behavior. However, the contribution of food information to the regulation of the SCN network remains controversial. Here, we identified the effect induced by time-restricted feeding (TRF) at dawn, but not at another time widow, inducing a robust and long-term shift in locomotor behavior and increased wakefulness. Comparing the oscillations of intracellular Ca^2+^ signals in the SCN GABAergic neurons of freely moving mice, before and after TRF, revealed significant activation of these neurons in dawn-TRF mice. Moreover, RNA-seq profiling in the dawn TRF-induced behavioral changes identified altered expressed genes involved in regulating extracellular exosome, ion transporters, and ECM-receptor interaction, but not core clock genes. Furthermore, injection in the SCN of insulin-like growth factor (IGF2) inhibitor Chromeceptin, targeting the most upregulated gene in extracellular exosome, abolished the after effect induced by ZT0-4 TRF. Finally, GABAergic-neuron-specific disruption of the potassium-chloride cotransporter *Kcc2* intensified the dawn TRF-induced after effect, indicating that *Kcc2* encodes food intake derived signals that control SCN clock entrainment. Thus, our study functionally links SCN GABAergic neuron activity and central clock entrainment regulation to both hunger- and food-response-related behaviors in mice.

## Introduction

Circadian rhythms are approximately 24-h cycles in physiology and behavior. The suprachiasmatic nucleus (SCN) is the central circadian pacemaker in mammals that receives photic and non-photic information, integrates time-related information from tissues, and then transmits timing information to cells and tissues to regulate physiology and behavior to entrain animals to the daily changes (Challet, 2019; Welsh et al., 2010). Also, the SCN encodes seasonal information that adjusts the onset and offset of activity depending on photoperiod length, notably, such changes can persist even under constant darkness (DD) (Farajnia et al., 2014; Houben et al., 2009; Inagaki et al., 2007; Kon et al., 2014; Olde Engberink et al., 2020; Schaap et al., 2003; VanderLeest et al., 2007). These results suggested that the SCN has a remarkable plasticity to adapt to environmental stimulation, with particular responsiveness to light.

Recent data collected for night shift workers, individuals with sleep disorders, and frequent flyer individuals (assessing jet-lag impacts) have provided evidence that metabolism and circadian rhythms are tightly linked (Challet, 2019; Huang et al., 2011). Feeding that follows a typical pattern of daytime eating for diurnal animals or nighttime eating for nocturnal animals affects metabolic homeostasis (Fonken & Nelson, 2014; Liu et al., 2014). A very recent study showed that feeding a piece of chocolate at the onset of the active phase prevented circadian desynchrony, whereas feeding chocolate near dawn prevented re-entrainment (Escobar et al., 2020).

The fact that mice given a 4-hour time-restricted feeding (TRF) at 21°C causes death, viewed alongside the observation that this phenotype can be rescued by impairing SCN function (Zhang et al., 2020), clearly suggests that the SCN somehow respond to TRF induced hunger-link food stimulation and regulate neuronal plasticity to modulate adaptive behaviors. The available evidence suggests that cyclic adenosine monophosphate (cAMP) is both an output of the SCN and an integral component of the SCN, regulating transcriptional cycles (Hastings et al., 2019; O’Neill et al., 2008). Also, dynamic regulation of SCN excitability appears to be tied to the redox state, apparently through nontranscriptional modulation of multiple potassium (K^+^) channels (Wang et al., 2012). Previous studies have also shown that restricted daytime feeding increases vasopressin release from the rat SCN (Aton et al., 2005), and resets the circadian clock in the SCN of white-footed mice and inbred CS strain house mice (Abe et al., 2007). Together, these findings implicate multiple factors as potential influencers of the rhythm-related functions of the SCN, including the TTFL, ion channels, neurotransmitters, neuropeptides, and/or the temporal variability of SCN neuronal coupling strength.

In the present study, we conducted unbiased screening to identify any entrained behavioral aftereffect(s) from time-restricted feeding. Our phase-response curve analyses indicated that mice are most responsive to TRF entrainment at dawn, and we carefully show that the delayed offset of activity from near dawn TRF persists in these animals, even after re-exposure to *ad libitum* feeding and 12:12 LD. Our data from long-term, fiber photometry monitoring of intracellular Ca^2+^ signaling in the SCN of freely moving mice revealed that the higher frequency spikes detected for the dawn TRF animals are mediated by the GCaMP7s Ca^2+^ signals, specifically in SCN GABAergic neurons. An RNA-seq analysis further showed that dawn TRF does not impact the transcription of core clock genes; rather, it deeply reprograms the expression of insulin-like growth factor 2 (IGF2), its high-affinity binding protein IGFBP6 and ion transporters. Blocking IGF2 abolishes dawn TRF induced locomotor aftereffect, and depletion of enriched *Kcc2* in the SCN GABAergic neurons exacerbates locomotor changes. Thus, our study links energy sensing and the ion physiology of particular SCN neurons to behaviorally consequential rhythmic entrainment by feeding.

## Results

### Time-restricted feeding entrainment generates persistent after effect in locomotor activity

To study the impacts of time-restricted feeding (TRF) on the circadian clock, C57BL/6J mice (6-8 weeks of age) were housed individually in cages with a wheel under a 12 h/12 h of light/dark (12:12 LD) cycle for two weeks, with free access to food (*ad libitum*). Then these mice were then subjected to two weeks of treatment with 1 of 12 different 4-hour TRF windows plus *ad libitum* feeding as control (n = 6 per group, with a 2-hour binning design). Afterwards, all mice were released into constant darkness (DD) and returned to *ad libitum* feeding (Fig. 1A). All tested mice showed that food intake gradually recovered (Fig. 1B), and body weights that were initially reduced but gradually recovered to the baseline level (Fig. 1C). We detected premeal activity (i.e., virtually the same as the “anticipation activity”) described by in (Fuller et al., 2009; Mistlberger et al., 2009) during the TRF phase of the experiment. However, such premeal activity disappeared soon after returning the animals to *ad libitum* feeding and release to DD (Fig. 1D).

**Fig. 1.**
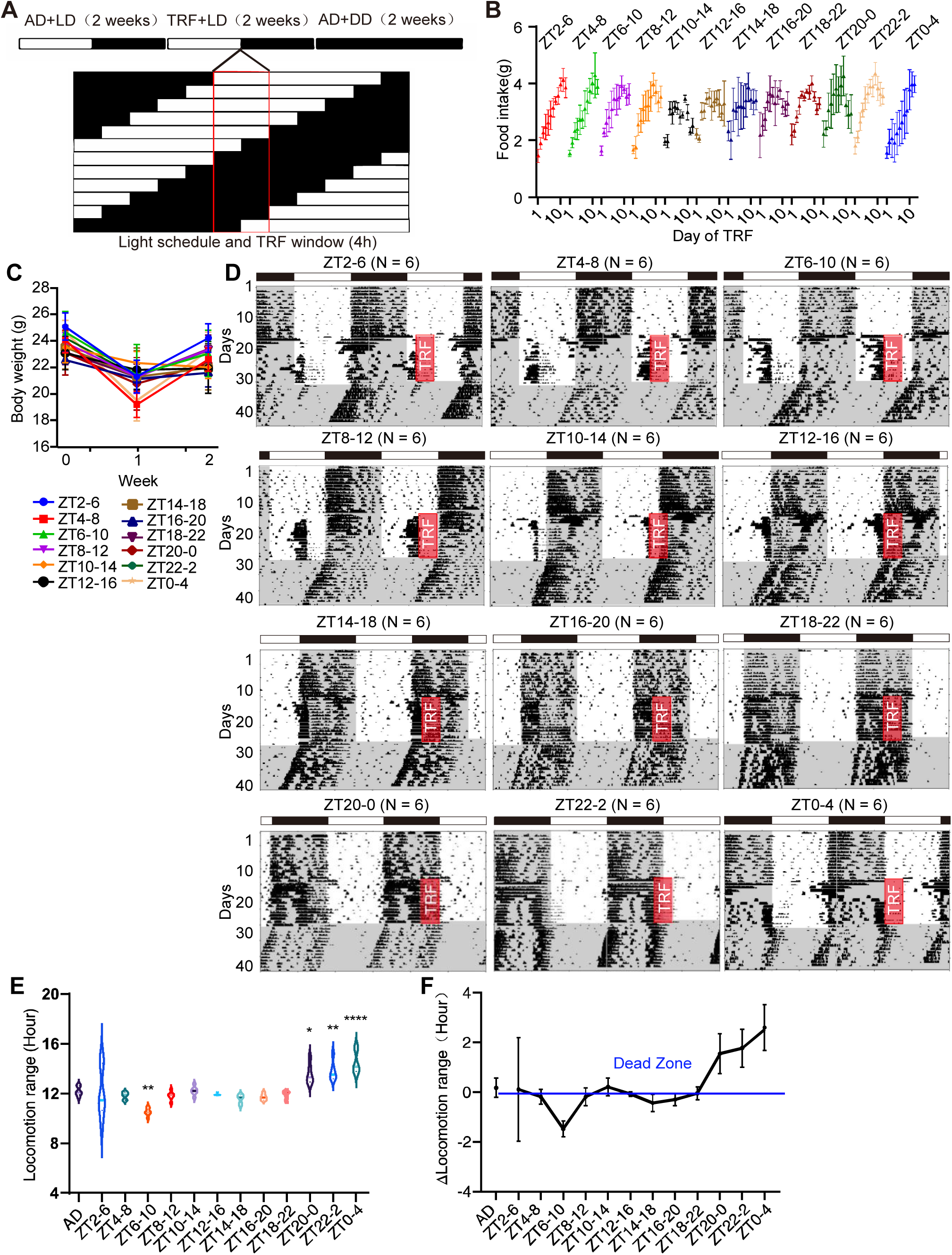
The effect of TRF on mouse locomotor. (**A**) The experimental design for time-restricted feeding. The time of light-dark cycle was changed over time to realize TRF at different phases. (AD: *ad libitum*, LD: Light-Dark cycles, DD: Dark-Dark cycles.) (**B**) The average of daily food intake from the first day of the TRF to 12th day in 12 phases TRF. (**C**) Weekly body weight of 12 phases TRF entrainment. 0 represents the week before the TRF (n = 6 each group). (**D**) The representative actogram of the TRF at 12 different phases (ZT2-6, ZT4-8, ZT6-10, ZT8-12, ZT10-14, ZT12-16, ZT14-18, ZT16-20, ZT18-22, ZT20-24, ZT22-2 and ZT0-4. Red rectangle: TRF window. The grey area represents dark phase. The numbers of mice in each time point was indicated in the top of graph thereafter. (**E**) The locomotion ranges of 12 phases TRF entrainment including *ad libitum* as control mice (n = 6). Values represent the average ± SD., *: p < 0.05, **: p <0.01, ****: p < 0.0001. (**F**) The phase responsive curve response by the TRF entrainment. The locomotion ranges subtracted by 12-h corresponds to the TRF time.

The most apparent trend from this experiment was that we observed an aftereffect of the TRF entrainment on delayed offset locomotor activity (i.e., the end point of activity). Note that this TRF aftereffect persisted for many weeks after the TRF portion of these experiments. It was evident that the impact of the ZT0-4 TRF window on mouse offset behavior was much stronger than for any other windows (Fig. 1D). Specifically, ZT0-4 TRF entrained mice displayed a significant delay of offset activity with increased distance between the onset (i.e., the start time of activity) and offset activity in DD. However, ZT0-4 TRF entrainment did not induce free-running period changes compared to that of *ad libitum* feeding using X^2^ periodogram analysis (23.69±0.06 vs 23.63±0.23, p > 0.05). Further, the kinetic change of the free-running period assessed by their onset activity and offset activity was not significantly different in the ZT0-4 TRF entrained mice (23.69±0.06vs 23.68±0.11, p > 0.05).

We also quantified locomotion range from onset activity through to the end of offset activity from the 1st day to the 10th day of post-TRF *ad libitum* feeding under DD to support the calculation of a phase responsive curve (PRC) for TRF time window and locomotion activity (Fig. 1E and 1F). These curves highlighted that the extent of the offset delay expands for the ZT20 to ZT0 TRF windows (Fig. 1E and 1F), and showed that an abrupt, unstable fluctuation followed this expansion at ZT2. These results were suggesting that ZT0 is apparently the most TRF-sensitive time point (around dawn-break). We detected only weak impacts on locomotor activity for the ZT8 to ZT18 TRF windows, suggesting that these time windows may be “dead zones” for any TRF-mediated impacts.

Having detected that the ZT0-4 TRF windows triggered the longest elongations of offset activity, we next determined how many days these TRF induced locomotor effects persisted. Mice were again exposed to a 2-week TRF entrainment (ZT0-4 TRF), but were then released into DD for a full month with *ad libitum* feeding. We found that TRF entrainment induced locomotor plasticity lasts for at least one month (Fig. 2A and 2B). Moreover, when mice were directly released into a 12:12 LD cycle, the aftereffect induced by ZT0-4 TRF entrainment again persisted (Fig. 2C and 2D), despite our expectation that this locomotor behavior change would be immediately suppressed by light. Pursuing this further, we applied saturating light pulses at CT0-CT1, but observed no significant phase advances for onset or offset activity by X^2^ periodogram analysis (Fig. 2E and 2F), suggesting that ZT0-4 TRF induced locomotor behavioral change may be independent of the well-studied transcription-translational feedback loop (TTFL) (Herzog et al., 2017; Takahashi, 2017; Welsh et al., 2010).

**Fig. 2.**
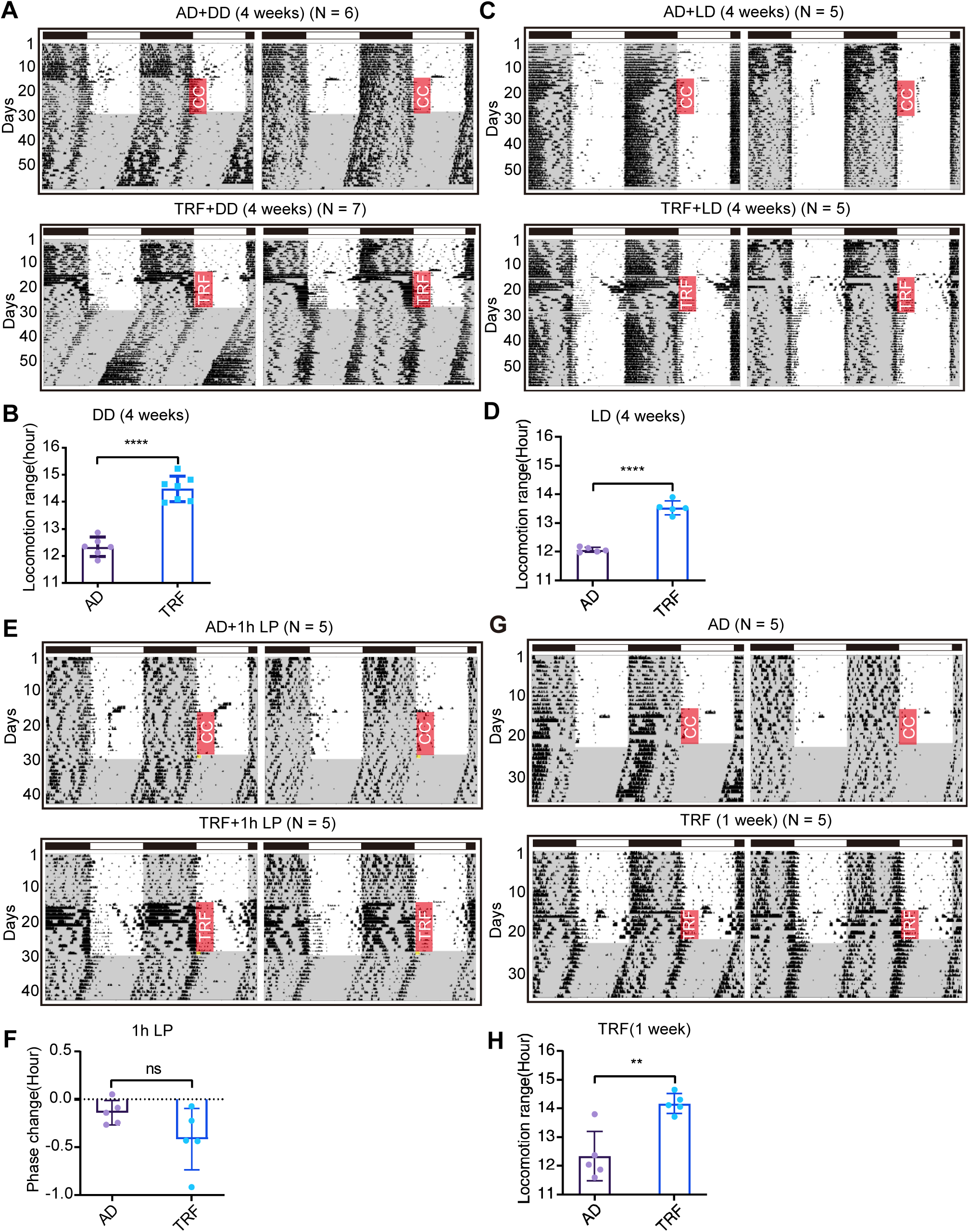
Robust aftereffect induced by iterative TRF. (**A**) The representative actogram of mice administrated with 2-weekTRF and released to DD and *ad libitum* for 4 weeks with *ad libitum* as control. CC means cages changes with food and thereafter. (**B**) Statistics of locomotion range under DD in the TRF and *ad libitum* group. (**C**) The representative actogram of mice administrated with 2-week TRF entrainment and released to LD and *ad libitum* for 4 weeks with *ad libitum* as control. (**D**) Statistics of locomotion range under DD in the TRF and *ad libitum* group. (**E**) The representative actogram of mice administrated with 2-week TRF and released to DD and *ad libitum* for 2 weeks with *ad libitum* as control. In the first day of DD, 200 lux light pulse (LP) is given at CT0-CT1 for 1 hour (yellow rectangle). (**F**) Statistics of phase shift under DD in the TRF and *ad libitum* group. (**G**) The representative actogram of mice administrated with 1-week TRF entrainment and released to DD and *ad libitum* for 2 weeks with *ad libitum* as control. (**H**) Statistics of locomotion range under DD in the TRF and *ad libitum* group. Values represent the average ± SD. **: p <0.01, ****: p < 0.0001, ns: not significant. Red rectangle: TRF window. The grey area represents dark phase.

We next conducted experiments in which we lesioned the dorsomedial hypothalamus (DMH), seeking to exclude potential effect from this region—which is known to mediate broad effects on mouse behavior and the expression of food-entrainable circadian rhythms (Fuller et al., 2008). Note that the SCN is known to project directly or indirectly into the DMH in a manner that impacts rhythmic control of food intake and locomotor behaviors (Saper et al., 2005). We found that lesion of the DMH did not abolish the locomotor plasticity induced by ZT0-4 TRF entrainment (Fig. S1), suggesting that DMH did not mediate the locomotor plasticity.

Finally, we investigated how shortening the TRF entrainment from 2 weeks to 1 week might affect this locomotor plasticity: under this condition, mice still exhibited a delayed offset activity, although the 1-week TRF entrainment induced a smaller effect than two-week TRF entrainment (Fig. 2G and 2H). These results firmly establish that restricting the feeding of mice to a window around dawn (ZT0-4) induces robust and persistent changes in locomotor behavior, and emphasize outsized impacts of this particular TRF window on mouse locomotor activity compared to TRF at other times of the day.

### The neuronal activation in the SCN of freely moving mice corresponds to the observed locomotor plasticity

Having identified that TRF induced locomotor plasticity which occurs each day and persists under DD or 12:12 LD, we then searched for the neuronal basis of this behavioral phenotype. Given the known role of the suprachiasmatic nucleus (SCN) in both circadian clock and output behavior (Hastings et al., 2018; Herzog et al., 2017; Inagaki et al., 2007; Mei et al., 2018; Shan et al., 2020), we initially focused our attention on this brain region. We made an effort to establish a recording system to monitor intracellular Ca^2+^ signals in the SCN of freely moving mice, specifically by using a jGCaMP7s, a high-performance (GFP-based) Ca^2+^ sensor that provides excellent detection resolution for individual spikes (Fig. S2A), a custom high-numerical aperture (0.5 NA) to increase signal-to-noise ratios (Fig. S2B), and a custom-designed dual-color optical fiber for photometry, with a Ca^2+^-dependent excitation wavelength at 470 nm and a Ca^2+^-independent isosbestic excitation wavelength at 410 nm (Fig. S2C).

After stereotactic injection of an adeno-associated virus (Alvarez-Saavedra et al.) encoding jGCaMP7s (together with AAV encoding Cre-dependent GABA transporter (VGAT) promoter) into the SCN (Fig. S2A and S2B) and allowing three weeks for full viral expression, we recorded Ca^2+^ signals originating from GABAergic neurons in the SCN of freely moving mice (Fig. S2D). The signal was recorded for 35s every 10 mins to reduce the phototoxicity to the SCN neurons using a custom Iper Studio Alpha script. Another custom-developed script was used to realize data splicing, as well as qualitative analysis of changes in different types of Ca^2+^ signals (spectral analysis) for the long-term recording dataset. Changes in fluorescence over baseline fluorescence (ΔF/F) were calculated as previously described (Jones et al., 2018). Before conducting any ZT0-4 TRF entrainment, we monitored *ad libitum* fed mice in constant darkness conditions from ZT12 to CT72.

We observed robust daily rhythms of intracellular Ca^2+^ spikes in SCN GABAergic neurons. The mean Ca^2+^ signals for SCN GABAergic neurons was almost 3× higher during the subjective daytime than at subjective nighttime (Fig. S2D). At dawn, Ca^2+^ signals showed a gradually ascending signals (around CT22 to CT24), while at dusk, Ca^2+^ signals fell quickly (around CT35 to CT37). The transition between the rising and the falling phases of the Ca^2+^ signals correspond separately to the end time offset of activity and onset of activity around dawn and dusk (Fig. S2D). The changes of Ca^2+^ signals reflect a real-time locomotor state, suggesting that Ca^2+^ signals enable us to dissect functional changes corresponding to the onset or offset of locomotor activity.

### TRF induces alteration of SCN GABA neuronal activity and their outputs

To assess whether TRF induced locomotor plasticity was related to patterns of SCN activity, we designed an experiment involving: a) recording of Ca^2+^ signals we performed over an 84-hour *ad libitum* feeding phase, b) a ZT0-4 TRF (including ZT8-ZT12 TRF for comparative analysis) entrainment for 2-weeks without recording signals, and finally, c) recording of Ca^2+^ signals upon release of these TRF exposed mice into DD for another 84-hour *ad libitum* feeding phase (Fig. 3A). We decided to abandon recording during the TRF entrainment phase because the mice’s excessive locomotor activity frequently broke the optical fibers. Moreover, the Ca^2+^ signals from TRF entrainment had substantially more inferior signal-to-noise resolution than the experiment’s initial and final phases.

**Fig. 3.**
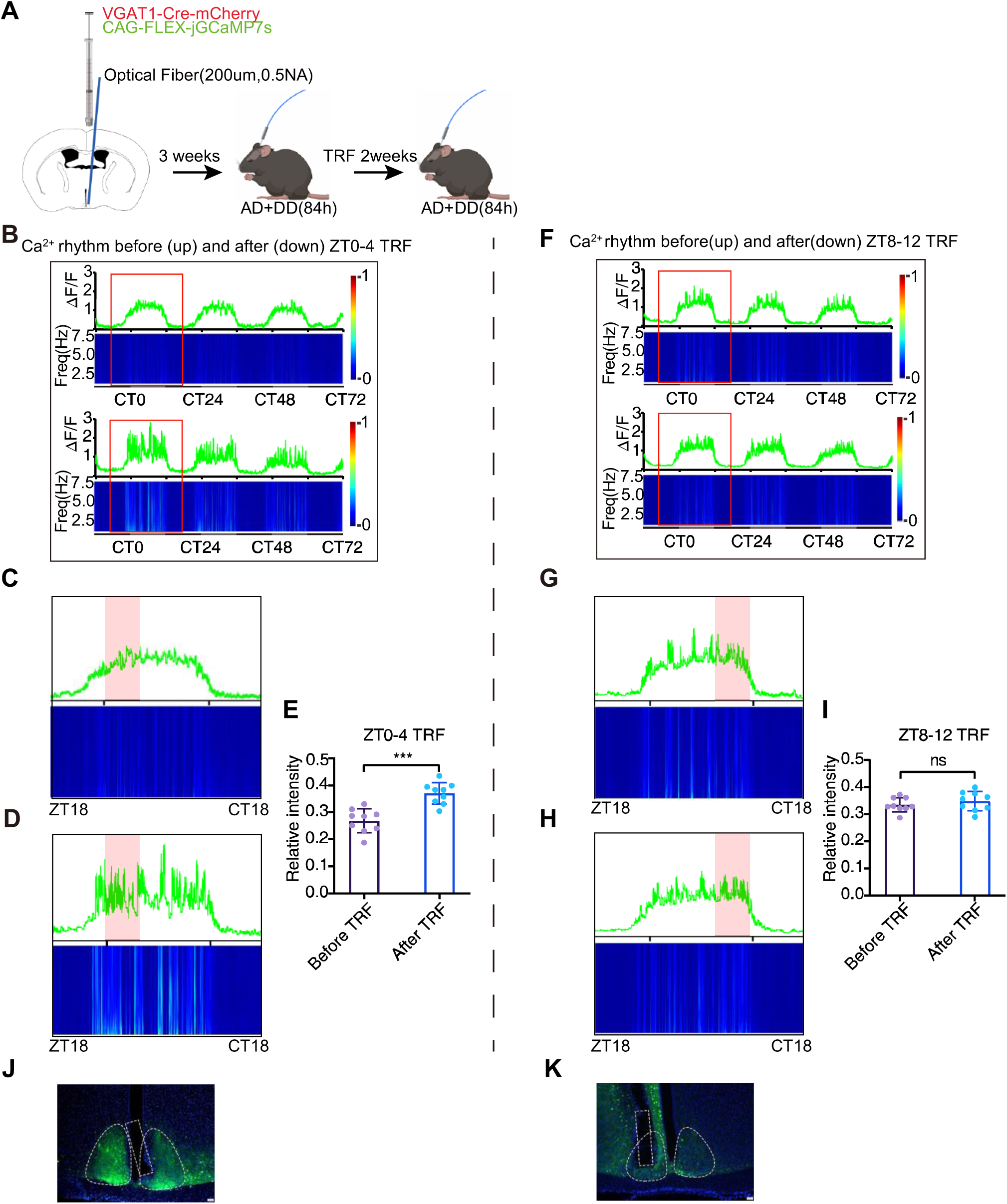
The elevated activities in the SCN GABAergic neuronal population by the TRF. (**A**) Experiment design of Ca^2+^ signal recording. The signal was collected before and after the TRF in the same mouse. (**B**) Ca^2+^ signals before (up) and after (down) TRF entrainment at ZT0-4. The heatmap shows the power of Ca^2+^ signal in different wave frequencies which is corresponding to the ΔF/F data. (**C-D**) A zoom out of Ca^2+^ signal from pink unfilled rectangle in Figure B. Pink solid rectangle: subjective TRF window before **c** and after **d**, i.e. *ad libitum*. (**E**) Relative intensity of Ca^2+^ signal before or after ZT0-4 TRF in each cycle (n = 9). The frequency power in 1 hour before subjective TRF window (ZT23-CT3, CT23-CT27, CT47-CT51) was normalized to total Ca^2+^ signal active phase (ZT23-CT13, CT23-CT37, CT47-CT61) Values represent the average ± SD. ***: p < 0.001. (**F-H**) The effect of the TRF entrainment at ZT8-ZT12 on Ca^2+^ signal recording as described in (**b-d**). (**I**) Relative intensity of Ca^2+^ signal before or after ZT8-12 TRF in each cycle (n = 9). The frequency power in 1 hour before subjective TRF window (CT7-CT11, CT31-CT35, CT55-CT59) was normalized to total Ca^2+^ signal active phase (ZT23-CT12, CT23-CT36, CT47-CT61)) Values represent the average ± SD. ns: not significant difference. (**J**) The expression of GCamp7s (Green) and DAPI (blue) and the location of optical fiber (white dotted box) in 50 μm SCN slice for ZT0-4 TRF mice. (Scale bar: 100 μm). (**K**) The expression of GCamp7s (Green) and DAPI (blue) and the location of optical fiber (white dotted box) in 50 μm SCN slice for ZT8-12 TRF mice. (Scale bar: 100 μm).

We observed a striking pattern in the frequencies of Ca^2+^ transients when we aligned the Ca^2+^ signals from the mice before and after ZT0-4 TRF: the Ca^2+^ transients for roughly 4-hour time windows each day were much more intense in mice that had been entrained with TRF (Fig. 3B). Specifically, the average Ca^2+^ transient frequency in such 4-hour windows was 0.36 ±0.04 (ranging from 1-7.5 Hz) in the ZT0-4 entrained mice but was only 0.27 ± 0.04 (ranging from 1-7.5 Hz) for the same mice before the ZT0-4 entrainment (Fig. 3C vs 3D, and quantitative analysis in Fig. 3E). Confirming the dawn TRF’s extraordinary impact, we detected no significant changes in Ca^2+^ transient frequency before and after the ZT8-12 TRF entrainment (Fig. 3F-H, and quantitative analysis in Fig. 3I). We confirmed the correct position of the fiber insertion in the SCN region and the expression of the GCaMP after each of above experiments, thus ensuring that the detected Ca^2+^ signals were indeed from the SCN (Fig. 3J and 3K). At a minimum, these observations indicate that TRF entrainment during a dawn time window encodes highly rhythmic patterns of differential SCN GABAergic neuron activity that persist for days after entrainment.

SCN^VIP^ neurons represent integration points for both photic and non-photic sensory inputs (Acosta-Galvan et al., 2011). The activation of SCN^VIP^ neurons by bilateral injections of Cre-dependent hM3Dq into the SCN of heterozygous VIP-Cre mice at ZT4 and ZT13 does not change locomotor or sleep-wake cycle phenotypes (Todd et al., 2020). Hence, we asked whether selective inhibition of SCN^VIP^ neurons may be able to simulate the effect of ZT0-4 TRF entrainment (i.e., delay the offset of activity). We injected AAV-DIO-hM4D(Gi)-mCherry into the SCN of VIP-Cre mice (n = 6) and administered the chemogenetic ligand clozapine N-oxide (CNO; 3 mg/kg) at ZT 0 for 13 consecutive days (Fig. S3A and S3B). We found that inhibition of SCN^VIP^ neurons during the daytime did not delay locomotor activity offset (Fig. S3C), indicating that simple activation or inhibition of SCN^VIP^ neurons during daytime cannot directly affect locomotor plasticity (Collins et al., 2020). Alternatively, SCN^VIP^ neurons may not involve in mediating TRF-related behaviors.

Given that TRF can entrain SCN GABAergic neuronal activity, we wondered whether ZT0-4 TRF would induce an aftereffect of sleep-wake cycles, one of the most important output of the SCN. We conducted EEG and EMG recording before and after the ZT0-4 TRF entrainment. The duration of non-rapid eye movement (NREM) and REM episode after ZT0-4 TRF entrainment were decreased substantially in the subjective TRF windows (CT0-4, CT24-28, and CT48-52) (Fig. 4A and 4B) compared to before TRF. Similar to increased locomotor activity, the wakefulness after ZT0-4 TRF entrainment was increased in the subjective TRF window (Fig. 4C and 4D). Thus, our data suggest that ZT0-4 TRF can entrain SCN output and induce persistent changes in sleep-wake cycles even withdraw TRF.

**Fig. 4.**
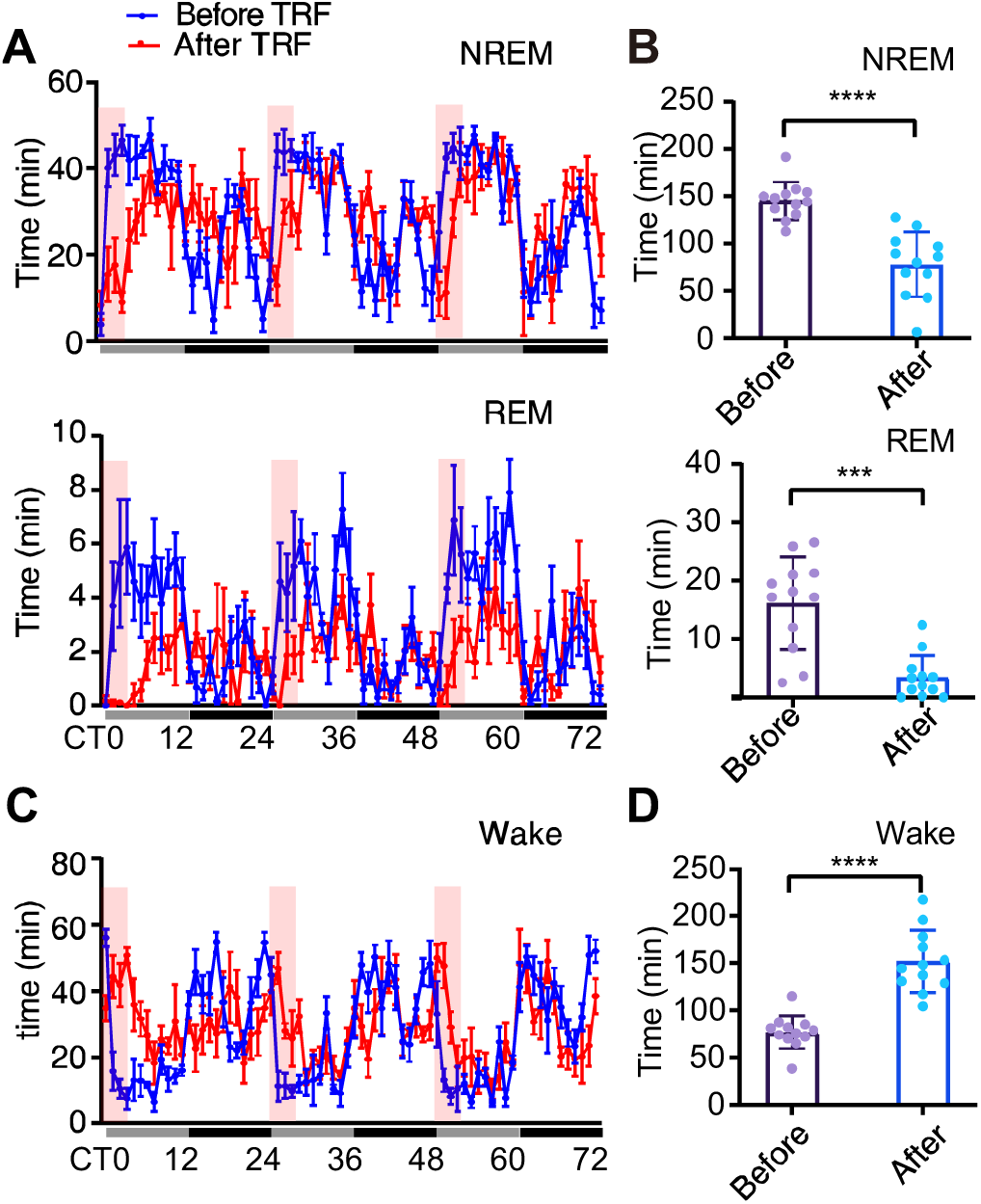
ZT0-4 TRF entrainment induces an after effect in the Sleep-wake cycle. (**A**) Changes in NREM sleep and REM sleep before and after ZT0-4 TRF under constant darkness. Values represent the average ± SEM, (n = 4 each group). Red rectangle: subjective TRF window. (**B**) Total time spent in NREM and REM in subjective TRF window (CT0-CT4, CT24-CT28, CT48-CT52) before or after TRF. Values represent the average ± SD. ***: p < 0.001, ****: p < 0.0001. (**C**) Changes in wake-up time before and after ZT0-4 TRF under constant darkness. Values represent the average ± SEM, (n = 4 each group). Red rectangle: subjective TRF window. (**D**) Total time spent in wakefulness in subjective TRF window (CT0-CT4, CT24-CT28, CT48-CT52) before or after TRF. Values represent the average ± SD. ***: p < 0.001, ****: p < 0.0001.

### ZT0-4 TRF impacts the expression of the ion transporters in the SCN

Given our finding that ZT0-4 TRF entrainment specifically induced altered SCN neuronal activity at dawn, we hypothesized that time-specific input information about feeding might induce changes in signaling pathway activity in the SCN; reprograming its SCN neuronal activity. We pursued this idea by performing RNA-seq based profiling of ZT0-4 SCN samples at CT1, CT7, CT13, and CT19 (CT1 corresponds to 1h after the subjective onset of daytime (near dawn) under constant-dark conditions). To identify dawn-only differentially expressed candidate genes, we included two additional sets of control samples: ZT8-12 TRF entrained SCN samples and *ad libitum* SCN samples at CT1, CT7, CT13, and CT19. The expression profiles of core clock genes (*Bmal1, Clock, Per1, Per2, Cry1, Cry2*) from the ZT0-4 TRF entrained SCN samples did not differ significantly from ZT8-12 and *ad libitum* groups (Fig. 5A and Fig S4), suggesting that the TTFL was not significantly affected. Note that we did detect a dampened amplitude for the expression of the secondary loop factor *Nr1d1* at CT7 and for the known TTFL output target *Dbp* at CT13 in the ZT0-4 TRF entrained animals (Fig S4).

**Fig. 5.**
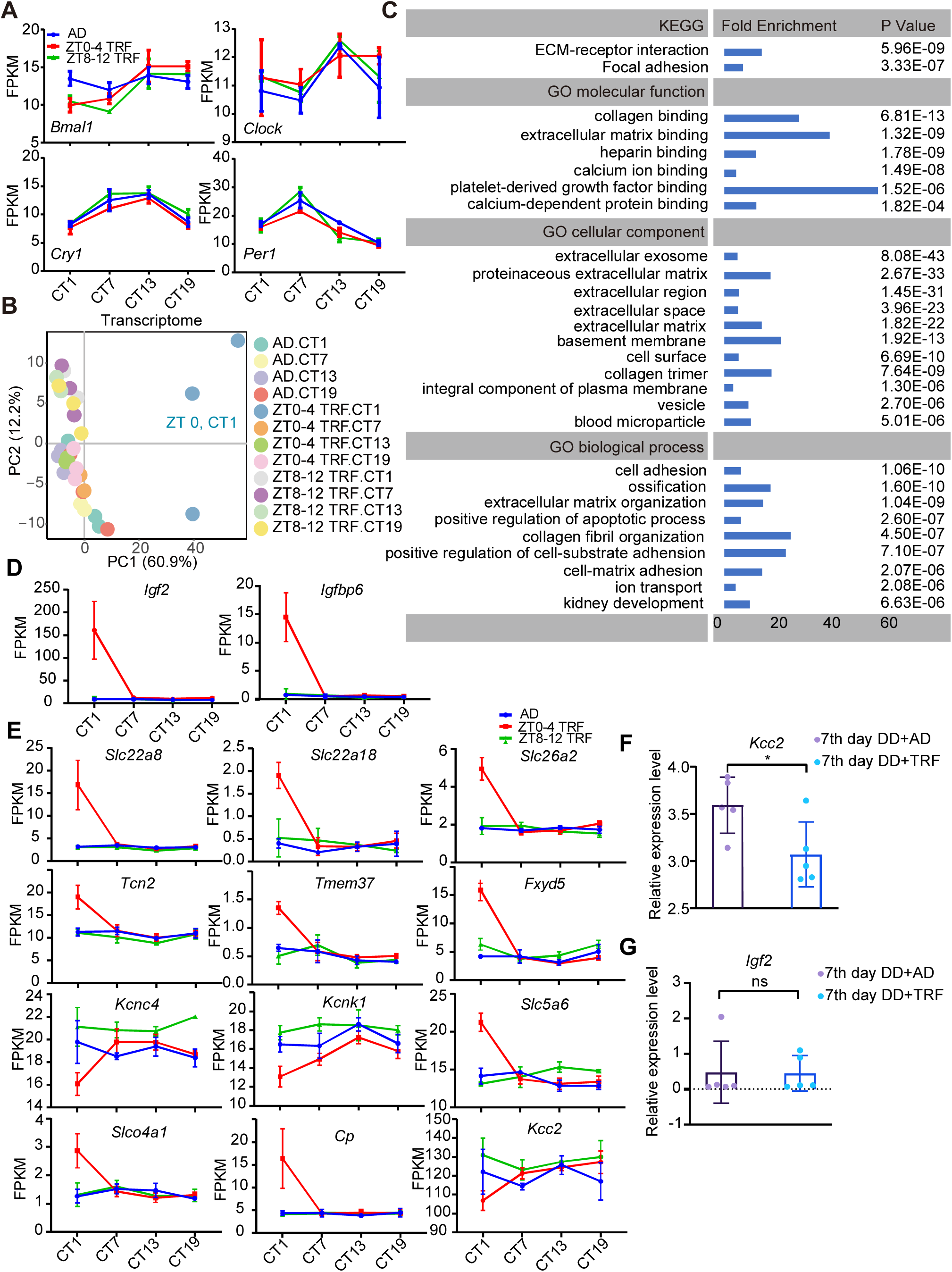
RNA seq profiling of the SCN samples showed the distinct transcriptome by the TRF entrainment at ZT0-4. (**A**) FPKM of clock genes in *ad libitum*, ZT0-4 TRF, and ZT8-12 TRF entrainment groups. Values represent the average ± SD. (**B**) PCA analysis of SCN RNA-seq raw data at CT1, CT7, CT13 and CT19 in three groups: ZT0-4 TRF, ZT8-12 TRF and *ad libitum*, (n = 3 per time point). (**C**) KEGG and GO pathway analysis in 142 most relevant genes. (**D**) Change of *Igf2* and *Igfbp6* in *ad libitum*, ZT0-4 TRF, and ZT8-12 TRF entrainment groups. Values represent the average ± SD. (**E**) Change of ion transport in *ad libitum*, ZT0-4 TRF, and ZT8-12 TRF entrainment groups. Values represent the average ± SD. (**F-G**) Relative expression level of *Kcc2* (F) and *Igf2* (G) in the CT0 SCN samples of the 7th day after ZT0-4 TRF under constant darkness in both *ad libitum* control and ZT0-4 TRF group. Values represent the average ± SD. *: p < 0.05.

Principal component analysis (PCA) was performed on the raw RNA-seq data (n =3 per time point) after Bartlett’s test. We saw that CT1 samples from ZT0-4 TRF (blue dots) were clearly distinct from the control samples (*ad libitum*), from the other ZT0-4 TRF or from ZT8-12 TRF, with evident separation along PC1 (Fig. 5B), suggesting that ZT0-4 TRF specifically reprograms the transcriptome of the SCN samples at CT1. Notably, this cluster corresponded to the observed high frequency of Ca^2+^ spike, and might therefore represent SCN neuronal activity. To analyze which transcripts exerted the most significant influence on PC1 in this model, we used the cos^2^ values to scale their relative importance (Nikhil et al., 2020). We shortlisted the top 214 genes that meet cos^2^ > 0.5, and interestingly found that 142 genes among 214 genes had significantly different expression in the ZT0-4 TRF entrained SCN at CT1 samples compared with other groups (See detailed in methods). Significant GO terms of the “KEGG” category included ECM-receptor interaction and Focal adhesion (Fig 5C). Significant GO terms of the “Molecular function” included collagen binding extracellular matrix binding, calcium binding etc. Significant GO terms of the “Cell component” included extracellular exosome, and the “Biological process” included ion transport (Fig 5C). Most of significantly differentially expressed genes (DEGs) in these enrichment pathways were related to various collagen proteins which have been reported to maintain structural and functional neuroplasticity by modulating Ca^2+^ and K^+^ channels and modulating intracellular Ca^2+^ concentrations (Fig. 5C) (Cho et al., 2018). In the secreted exosomes pathway, *Igf2* and *Igfbp6* are the most upregulated genes that may affect nutrient sensing and regulate neuronal plasticity to modulate adaptive behaviors involved in food seeking in the previous studies (Fig 5D)(Fernandez & Torres-Aleman, 2012).

We found that ZT0-4 TRF entrainment caused reprogramming of genes for ion transports including *Kcc2, Slc26a2, Slc13a3, Slc5a6, Slc22a18, Slc26a7, Slc22a8, Slc04a1, Kcnc4, Kcnk1, Tspo, Fxyd5, Wnk4, Cp, Cybb, Tmem37, Tcn2, Clic1* (Fig. 5E). We were particularly interested in *Kcc2* which was highest enriched among all altered ion transporters in the SCN and encoded a potassium-chloride co-transporter protein previously reported to function in a long day (i.e., seasonal) clock entrainment (Olde Engberink et al., 2018). To test whether the down regulation of *Kcc2* persists, the expression of *Kcc2* was examined using the SCN samples from 7^th^ day after releasing to *ad libitum* in experienced ZT0-4 TRF mice. We found that the level of *Kcc2* expression significantly decreased in TRF SCN compared with *ad libitum* SCN (Fig 5F), indicating that the state of *Kcc2* coincides with the changes of TRF-induced long-term locomotion. Interestedly, the expression levels of *Igf2* and *Igfbp6* quickly recovered to the normal levels after withdrawing TRF (Fig 5G), suggesting that *Igf2* and *Igfbp6* sense energy homeostasis and coordinate behavioral in the SCN.

Ion channels have been previously demonstrated as significant regulators of functional SCN pacemaker formation, including responses to photoperiod changes (Choi et al., 2008; Farajnia et al., 2014; Meredith et al., 2006; Montgomery et al., 2013; Myung et al., 2015). There are obvious similarities between the previously reported seasonal-change-induced phenotypes upon *Kcc2* disruption and our observations of locomotor plasticity induced by TRF: both our ZT0-4 TRF entrainment and long-day entrained mice (Rohr et al., 2019) showed decreased *Kcc2* levels and displayed altered locomotor length.

### Kcc2 encodes food intake derived signals in the SCN that control locomotor outputs

Considering TRF induced the reprogram of ion transporters, and that *Kcc2* is the most abundantly expressed in the SCN among these ion transporters, we knockdown the *Kcc2* in the SCN GABAergic neurons by stereotactically injecting AAV-VGAT1-Cre and AAV-Ef1a-mCherry-U6-Loxp-CMV-EGFP-loxp-shRNA (*Kcc2*) or scramble shRNA into the SCN, enabling *Vgat1Cre*-dependent expression of the shRNA. Locomotor activity was monitored (Fig. 6A and 6B). The expression of KCC2 and mCherry in all mice was examined by immunostaining or fluorescence detection after the experiment to ensure that *Kcc2* was specifically knocked down in *Vgat1* neurons.

**Fig. 6.**
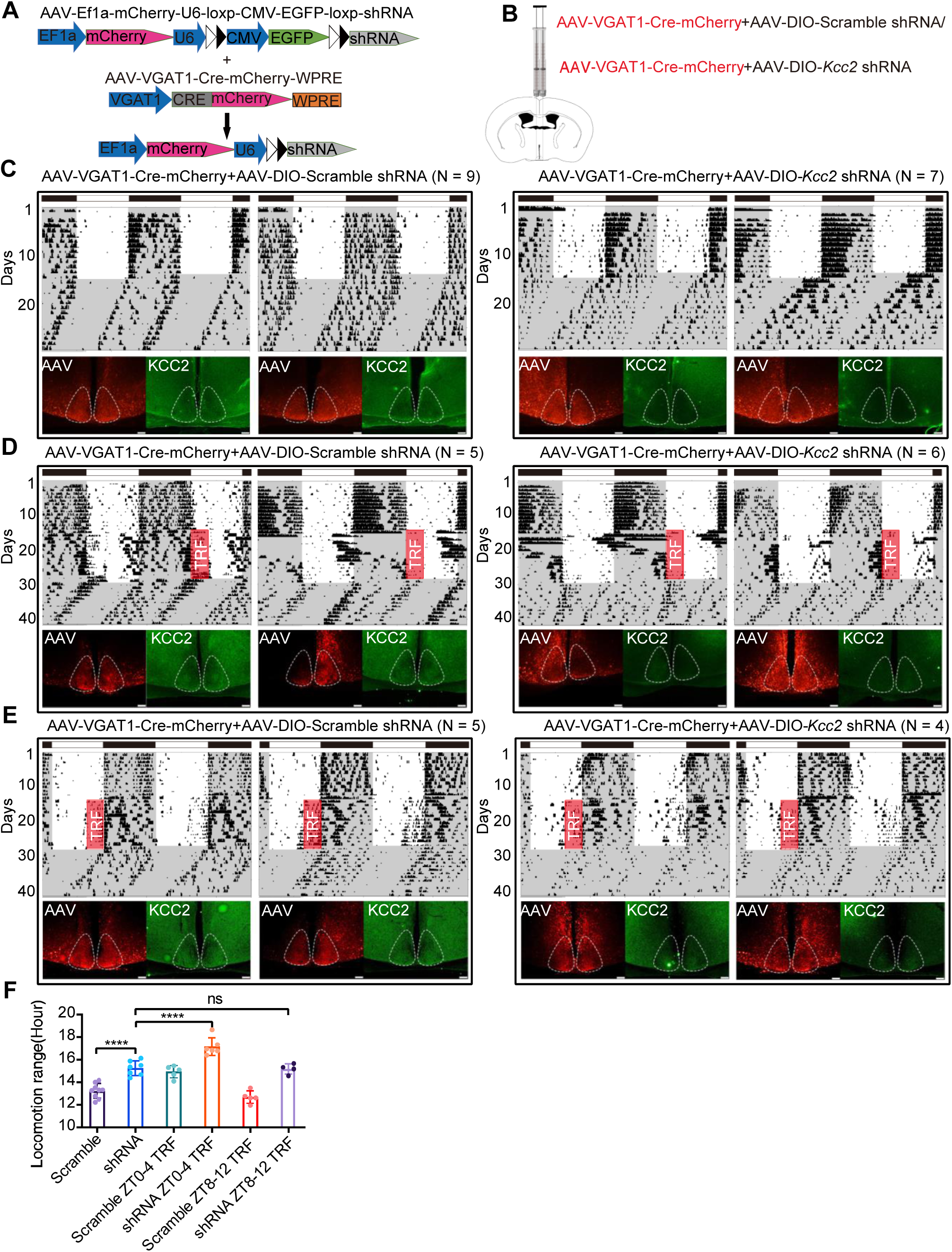
Depletion of *Kcc2* in the SCN GABAergic neurons exacerbates the locomotor plasticity in response to the ZT0-4 TRF. (**A**) The skeleton of AAV-VGAT1-Cre and Cre inducible AAV-shRNA. (**B**) A mixture of AAV-Cre and AAV-*Kcc2* shRNA or AAV-scramble (150 nl) were injected in SCN bilaterally. (**C**) The representative actogram of locomotor in control and *Kcc2* knockdown mice. All mice were entrained under LD for 2 weeks and then released to DD for 2 weeks. The expression of KCC2 (Green) was verified by anti-KCC2 in SCN slice after experiments. The AAV injection site (Red) was also verified by SCN slice. (Scale bar: 100 μm) (**D**) The representative actograms of locomotor in the control and *Kcc2* knockdown mice after 2 weeks ZT0-4 TRF entrainment. (Scale bar: 100 μm) (**E**) The representative actograms of locomotor in the control and *Kcc2* knockdown mice after 2 weeks ZT8-12 TRF entrainment. (Scale bar: 100 μm) (**F**) Statistics of the locomotion range of each group. Values represent the average ± SD. ****: p < 0.0001, ns: not significant difference.

Behavioral assays with these animals revealed that knockdown of *Kcc2* in the GABAergic neurons of SCN significantly increased the overall locomotion range between forward onset activity and backward offset activity (Fig. 6C). Because the circadian oscillation in the posterior SCN was phase-locked to the offset of activity, while the anterior SCN was phase-locked to the onset of activity (Inagaki et al., 2007), this apparent decompression of the onset and offset of daily activity is most likely a consequence of reduced posterior SCN neuronal and anterior SCN neuronal coupling strength. *Kcc2* is known to mediate SCN coupling, and a previous study demonstrated that KCC2 encodes seasonal plasticity information (Myung et al., 2015; Rohr et al., 2019). We found that *Kcc2* knockdown mice entrained with ZT0-4 TRF displayed larger dispersions of onset and offset activity than *ad libitum* feeding *Kcc2* knockdown mice (Fig. 6D). In contrast, no such additive phenotype was observed with corresponding experiments for ZT8-12 TRF entrainment (Fig. 6E and 6F). Our results show that depletion of *Kcc2* in SCN GABAergic neurons further exacerbates TRF entrained locomotor plasticity, supporting the notion that the KCC2 protein somehow encodes food intake information in the SCN.

To test whether TRF interferes with light entrainment, mice were entrained under long photoperiod (16:8 LD) for 2-weeks, and then given ZT0-4 TRF entrainment for further 2-weeks (Fig. 7A). These experiments included control mice (12:12 LD), and activity onset and offset times were measured daily after releasing them into DD. We detected no differences in the average daily onset times for activity among 16:8 LD entrained mice, LD plus ZT0-4 TRF entrained mice, or the 12:12 LD mice (Fig. 7B). In striking contrast, there was a clear difference in offset times among these entrainment groups. Specifically, the 16:8 LD plus ZT0-4 TRF entrainment mice displayed the same dynamic changes in offset time as the 12:12 LD mice, whereas the 16:8 LD entrained mice without TRF entrainment took fully 7 days to recover to the offset times of the 12:12 LD control mice (Fig. 7B). Furthermore, *ad libitum* mice were hardly entrained to short photoperiod (8:16 LD), while ZT0-4 TRF mice showed significantly delayed offset to match to light-off time, suggesting that ZT0 TRF could facilitate the adaptation of the central clock to a long photoperiod (Fig 7C and 7D).

**Fig. 7.**
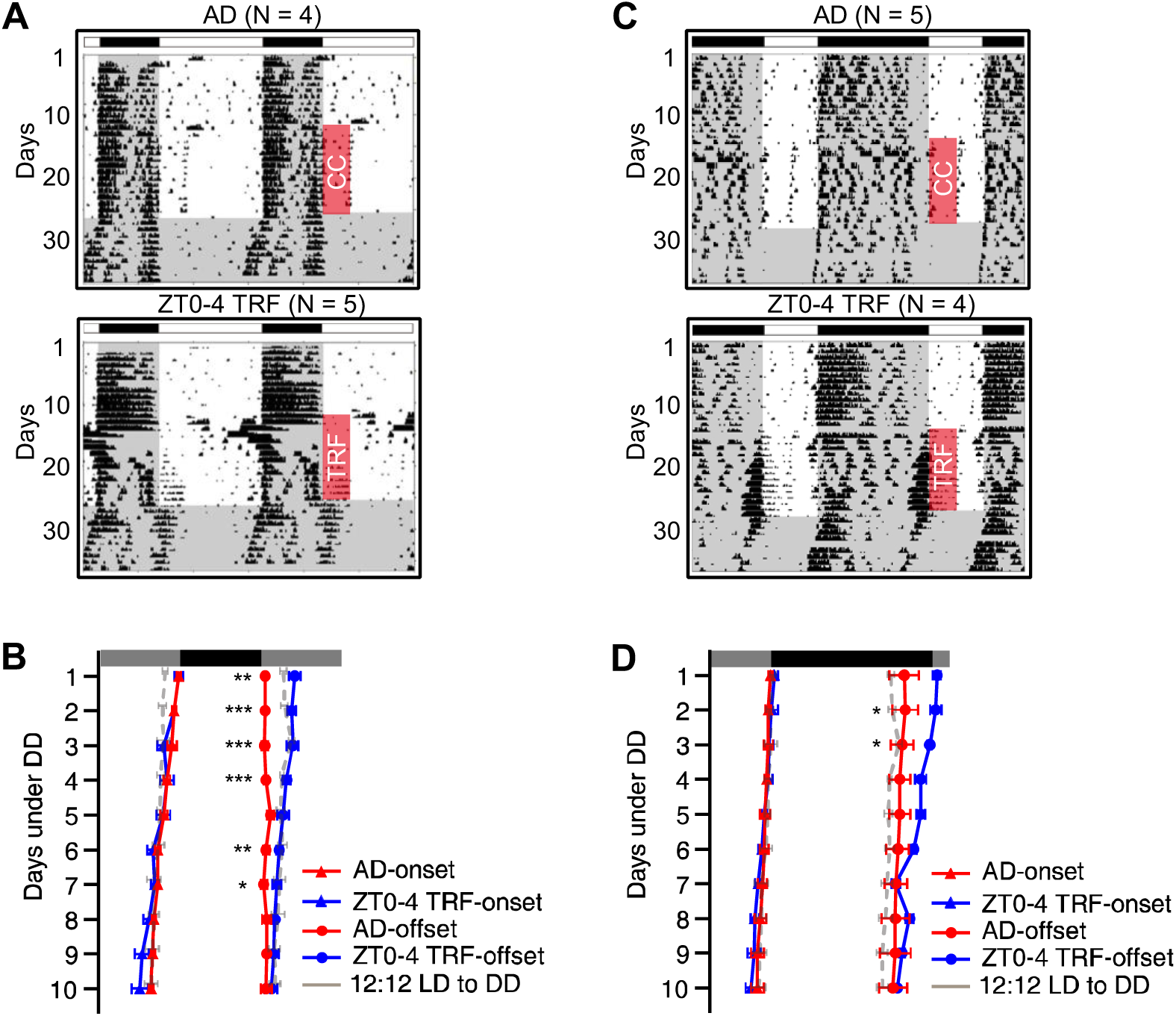
ZT0-4 TRF entrains central clock in both long photoperiod and short photoperiod. (**A**) Representative actograms of locomotor in the long days (16:8 LD) w/o TRF ZT0-4. (**B**) The dynamic adaption of onset and offset activity in the DD after ZT0-4 TRF entrainment and long days entrainment. Grey lines represent the recovery curves from 12:12 LD entrainment mice. Values represent the average ± SD. **: p <0.01, ***: p < 0.001. (**C**) Representative actograms of locomotor in the short days (8:16 LD) w/o TRF ZT0-4. (**D**) The dynamic adaption of onset and offset activity in the DD after ZT0-4 TRF entrainment and short days entrainment. Grey lines represent the recovery curves from 12:12 LD entrainment mice. Values represent the average ± SD. *: p <0.05.

These intriguing findings indicate that dawn TRF entrainment (ZT0-4) can interfere with the photoperiod entrainment process.

### Administration of IGF2R inhibitor (Chromeceptin) in the SCN blocks ZT0-4 TRF induced locomotor changes

The potential cooperative action of *Igf2* and its high affinity binding protein *Igfbp6* together with ion transporters led to investigate whether inhibition of IGF pathway blocks ZT0-4 TRF induced locomotor changes. A stainless steel brain cannula (RWD brain infusion kit) connected to the pump was inserted in the SCN, and IGF2R inhibitor-Chromeceptin or IGF1R inhibitor-GSK1904529A were administrated for two weeks during ZT0-4 TRF. Notably, inhibition of IGF2R completely abolished the aftereffect induced by ZT0-4 TRF (Fig 8A and 8B), suggesting that the observation of elevated *Igf2* and *Igfbp6* in the ZT0-4 TRF treated mice is responsible for delayed locomotor offset. In line with no change in the expression of *Igf1*, IGF1R inhibitor did not interfere delayed locomotor offset, indicating that *Igf2* and *Igfbp6* represent a specific energy sensor in the SCN. Altogether, this evidence indicates that the elevation of *Igf2* and *Igfbp6* levels in the SCN under ZT0-4 TRF mediates the aftereffect of locomotor. Considering that the levels of *Igf2* and *Igfbp6* quickly reversed to normal level after withdrawing TRF (Fig 5), while downregulation of *Kcc2* persisted, we proposed that *Kcc2* is downstream of IGF2-IGFBP6 pathway which coordinate locomotor adaptation to food seeking with long-term effect (Fig 8C).

**Fig. 8.**
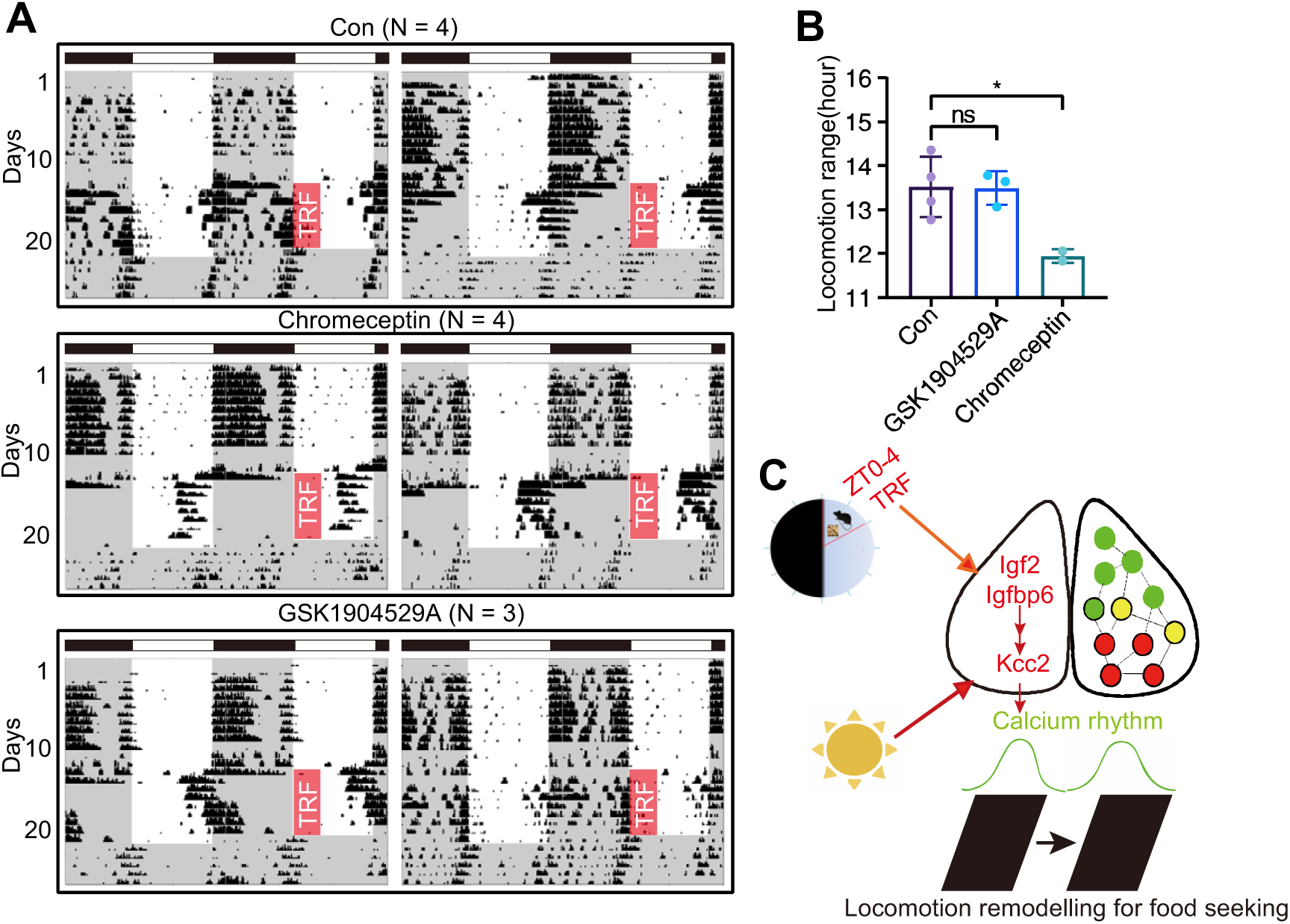
IGF2R inhibition block the after effect of ZT0-4 TRF on locomotion change. (**A**) The representative actogram of control, IGF2R inhibitor-Chromeceptin and IGF1R inhibitor-GSK1904529A treated mice administrated with 1-week TRF entrainment and released to DD and *ad libitum*. (**B**) Statistics of locomotion range under DD in control, GSK1904529A and Chromeceptin treated mice. Values represent the average ± SD. *: p <0.05, ns: not significant. Red rectangle: TRF window. The grey area represents dark phase. (**C**) The working model of ZT0-4 TRF on SCN. ZT0-4 TRF induces up-regulation of IGF2 signaling and thereby affects the expression of ion transport such as *Kcc2,* and finally influences the circadian output including the neuron Ca^2+^ rhythm, sleep-wake cycles and locomotion.

## Discussion

This study found that iterative ZT0-4 TRF results in a post-dawn locomotor phenomenon and affects the sleep-awake cycle that manifesting as persistent behavioral change. Both Ca^2+^ signal recording of the SCN in freely moving mice and RNA-seq analysis of samples from TRF entrained and control SCN samples indicated that the post-dawn locomotor phenomenon is mediated by IGF2 pathway coordinated with ion transport dynamics, specifically in SCN GABAergic neurons. We found that blocking IGF2 pathway counteracted the aftereffect induced by ZT0-4 TRF. Moreover, selective disruption of the *Kcc2* ion channel in SCN GABAergic neurons exacerbates dawn TRF entrained locomotor changes. Thus, our study demonstrates that the SCN receives food intake signaling and consequently forms a persistent aftereffect, thereby functionally linking SCN GABAergic neuronal activity to the entrainment of both hunger food-response-related behavior in mice.

### Consequences from misaligned circadian rhythms and food intake

In humans, many dawn circadian phenomena have been demonstrated, including early-morning elevations of blood glucose and hormones (e.g., cortisol), as well as early-morning activation of autonomic nervous system activity (Ando et al., 2016; Oster et al., 2006). Medical implications of these exaggerations of everyday circadian phenomena include poor metabolic control on diabetes mellitus patients (Porcellati et al., 2013) and increased incidence of cardiovascular diseases in the morning (Crnko et al., 2019). The present study shows that dawn (ZT0-4) TRF entrainment induces a robust aftereffect on the extended locomotor range. Given our demonstration that this aftereffect originates from GABAergic neurons in the SCN, and considering the well-established similarity in the rhythmic activities of these neurons between nocturnal and diurnal mammals (Kuhlman, 2007), we infer that the TRF-induced dawn phenomenon observed in mice (a nocturnal species) cannot be interpreted as a simple predinner effect (dusk phenomenon).

Our data indicate that the persistent behavioral changes induced by near-dawn TRF resemble changes in locomotor activity as induced by seasonal photoperiod (Farajnia et al., 2014; Kon et al., 2014; Myung et al., 2015; Rohr et al., 2019). We show that the observed locomotor plasticity associates with *Kcc2* downregulation in the GABAergic neurons of the SCN of TRF ZT0-4 entrained mice and resembles the decreased *Kcc2* levels observed in the SCN that receives and processes light input (Rohr et al., 2019). At a minimum, this resemblance suggests that downregulation of *Kcc2* is somehow involved in SCN network recovery and associates with locomotor behaviors after ZT0-4 TRF and after exposure to long-day conditions. It seems likely that a *Kcc2* related mechanism may represent an adaption for food-seeking behavior during seasonal changes. However, we acknowledge that TRF-induced the reprogramming of ion transporters includes more than a dozen genes, and it is undoubtedly not only *Kcc2*.

Consequently, the network of SCN may represent a sensitive time window (at dawn) that resists input signals. It is known from shift-workers and from frequent airline travelers (jet-lag) that “internal desynchrony” can occur between internal rhythms and environmental cues (Bass & Takahashi, 2010), which can manifest as increased susceptibility to obesity, metabolic diseases, cardiovascular events, and cancer (Zarrinpar et al., 2016). Previous studies with a jet-lag model, restricting feeding to the “active phase” accelerates resynchronization, whereas restricting feeding to the rest phase exerts a disruptive influence on circadian synchronization (Guerrero-Vargas et al., 2018). A very recent study showed that feeding a piece of chocolate at the onset of the active phase prevented circadian desynchrony, whereas feeding chocolate near dawn prevented re-entrainment (Escobar et al., 2020). In the present study, our screening of various TRF time windows revealed that ZT0-4 TRF entrainment induced an aftereffect on locomotor plasticity and decreased sleep time near dawn, which coincides with the activation of the SCN GABAergic neurons. Of note, we found that the DMH ablation—a potential food-entrainable oscillator—did not abolish this dawn phenomenon, further supporting that the SCN can be directly entrained by food intake. Furthermore, the fact that activation of *Igf2* and *Igfbp6* in the ZT0-4 TRF and inhibition of IGFR2 completely abolished the aftereffect by ZT0-4 TRF provided conclusively evidence that feeding signaling can input to the SCN, consequently SCN coordinates behavior adaptation through reprograming ion transporter to food seeking. Whether IGF2 and IGFBP6 impact directly the ion-transporters or mediated by ECM-receptors in the SCN is presently unknown. Thus, further investigation of the mechanism(s) and factors through which TRF impacts SCN neuronal activity in particular time windows should help clarify the consequences of misaligned circadian rhythms and food intake.

### Entrainment of the SCN by food intake

SCN neurons support precise and robust timekeeping, and light is known to be a primary entraining trigger for the SCN. Photic information directly conveys to the SCN, which then regulates transcriptional and translation of clock genes (e.g., *Per1*) to pattern suitable output responses suitable to environmental changes (Hastings et al., 2018; Herzog et al., 2017). There is evidence to support that locomotor outputs from the SCN are only rarely affected by restricted feeding. Consistent with this, daytime TRF does not alter the expression of clock genes in the SCN of 12:12 LD-exposed mice (Damiola et al., 2000). When rats and C57BL mice under LD are fed a hypocaloric diet, significant phase advances of locomotor activity are observed.

Our present study provides direct evidence, obtained with fiber photometry recording of Ca^2+^ signals that intracellular Ca^2+^ signals of the SCN are responsive to food intake, with the most significance responsive window occurring at ZT0-4. This response was also evident in the feeding response curve data. It bears strong emphasis that this higher frequency of Ca^2+^ spikes in the ZT0-4 TRF entrained mice persists, even for *ad libitum* fed mice in DD and LD photo conditions. We interpret this to indicate that a learning component must be involved in this phenomenon, but any underlying mechanism(s) remain unknown. There is extensive evidence that memory formation is strongly affected by the time window of training, so identifying how this stable state is entrained in the SCN network should be a highly informative area of future work.

Recall our finding that dawn TRF-induced changes in Ca^2+^ signals in the SCN GABAergic neurons did not rephase or otherwise affect the expression of the core clock genes *Per1* or *Per2*. This result is also supported by our findings that the delayed offset of locomotor activity cannot be masked by light or shifted by light pulses. A previous study demonstrated that K^+^ leak currents are regulated by metabolic oscillation in SCN neurons, thereby illustrating a non-transcriptional mechanism through which the clockwork machinery can modulate membrane excitability (Wang et al., 2012).

Our study offers another demonstration for non-transcriptional mechanisms can mediate rhythmic changes in SCN neuronal activity (i.e., ZT0-4 TRF might cause the most hunger than any other time, activates the IGF2-IGFBP6 pathway and consequently modulate the ion transporter pathway, thereby coordinating locomotor adaptation to food seeking). Notably, IGF6 and IGFBP6 quickly returned to normal after TRF, but IGF-mediated changes in ion transport persists and changes SCN neuron plasticity, leading to aftereffects. We provide evidence that ZT0-4 TRF can apparently “compete” with long day entrainment in terms of locomotor outputs. Viewed alongside our data showing enrichment for ion transport pathways among the differentially regulated genes of the ZT0-4 entrained SCN samples, these results indicated that food intake/nutrition-related ion changes somehow entrain the SCN and attendant behavioral outputs and affect wakefulness. Thus, future investigations about how ZT0-4 TRF impacts ion transport in the SCN could reveal how food intake timing of impacts circadian rhythm entrainment.

## Materials and Methods

### Animals

C57BL/6J mice were housed in pathogen-free animal facilities. All animal procedures were approved by the Animal Care and Use Committee of Soochow University. Mice were fed a normal chow diet (ShooBree SPF Mice Diet, 28% protein, 13% fat, 57% carbohydrates). VIP-Cre mice are obtained by Soochow University.

### Feeding schedule, food intake, and locomotor activity analyses

Six to eight-week-old male C57BL/6J mice were individually housed in cages equipped with running wheels for 2 weeks under light-dark cycle, with unrestricted access to food and water. After two weeks, mice were subjected to TRF for two weeks. Mice from each experimental group subjected to TRF were given access to food for 4-h at 12 different phase (ZT0-4, ZT2-6, ZT4-8, ZT6-10, ZT8-12, ZT10-14, ZT12-16, ZT14-18, ZT16-20, ZT18-22, ZT20-24, ZT22-2, where ZT0 = lights on and ZT12 = lights off, n = 6). Cages were changed daily during TRF to avoid the residual food in the cage. *Ad libitum* control mice (n = 6) would undergo the same procedures but with food available all the time. Daily food intake was measured manually and body weight was measured weekly. Wheel running was recorded and analyzed using ClockLab (Actimetrics, Evanston, IL). The onset and offset of locomotion are recognized by ClockLab, and the locomotion range is calculated by average the distance of first 10 days after release to *ad libitum* under DD.

### Stereotaxic viral injections

AAV-hsyn-Cre and AAV-FLEX-taCasp3 were obtained from OBiO, Shanghai. AAV-CAG-FLEX-jGCaMP7s, AAV-VGAT1-Cre-mCherry, AAV-Ef1α-DIO-hM4D(Gi)-mCherry, AAV-Ef1α-DIO-mCherry, AAV-Ef1a-mCherry-U6-Loxp-CMV-EGFP-loxp-shRNA (Scramble) and AAV-Ef1a-mCherry-U6-Loxp-CMV-EGFP-loxp-shRNA (*Kcc2*) were obtained from the Brain VTA, Wuhan, China. In all cases, 2-3 months old animals were initially anesthetized in the isoflurane induction box (2.5% isoflurane) and placed in a stereotaxic head frame during surgery with 1.5% isoflurane. A heating pad was used to prevent hypothermia. Virus injections (150 nl for SCN and 100nl for DMH) were delivered with a Shanghai Gaoge Industrial 10 μL syringe at 100 nl/min using an injection pump (ALC-IP, Shanghai Alcott biotech co. ltd). Stereotaxic coordinates of SCN: AP −0.05 mm, ML ± 0.15 mm, DV −5.85 mm. Stereotaxic coordinates of DMH: AP −1.4 mm, ML ± 0.25 mm, DV −5.2 mm.

### Ca^2+^ signal recording and signal analysis

200 μm diameter optic fiber was implanted with 5 degree in SCN just after the AAV injection. After 3 weeks, all implanted mice were screened for the rhythmic Ca^2+^ signal using a multichannel fiber photometry. Then, freely-moving mice will be recorded at ZT12 (light off) and last for 3 days in constant darkness. After mice were entrained in LD for 3 days, subjected to TRF for 2 weeks and then were released into DD with *ad libitum*, Ca^2+^ signals were re-recorded at ZT12 (light off) as before TRF entrainment. The sampling rate is 30 Hz for 470 nm light as signal channel and 430 nm light as negative control channel with power of 15-20 μw. The signal was recorded 35S every 10 min to prevent the phototoxicity to SCN neurons. The 430 nm signal is used to eliminate the noise caused by mouse movement or environmental interruption. Then, the raw data was normalized to 0-100 to counteract the difference of fluorescence strength result from the deviation of optic fiber location and virus expression. We then use wavelet analysis to calculate time-series signal power with different frequency and compare the signal difference in 1-hour temporal resolution before and after TRF entrainment in both ZT0 TRF group and ZT8 TRF group.

### Sleep-wake monitoring and analysis

2-month-old mice were surgically implanted with HD-X02 biotelemetry transmitters (Data Sciences International [DSI], New Brighton, MN) according to the manufacturer’s protocol (DSI EET Device Surgery Manual). Mice were deeply anesthetized with isoflurane (2.5% induction, 1.5% maintenance) and immobilized in a stereotaxic apparatus. After exposing and cleaning the skull, two stainless steel screws were inserted above the dura membrane worked as cortical electrodes. The other pair of screws were inserted into the cervical trapezius muscles to record EMG. The electrodes above the skull were fixed with dental acrylic, and the transmitter was pocked under the back. Mice were allowed to recover for at least 7 days before starting the experiment. The EEG and EMG data were collected for 3 days from CT0 to CT72 for both before TRF and after ZT0-4 TRF.

EEG (filtered by 0-30 Hz) and EMG signal sampled at 500 Hz were analyzed using the Neuroscore version 3.3 software (Data Sciences International [DSI], New Brighton, MN). The delta (0.5–4 Hz) (NREM), alpha (8–12 Hz) (wake) and theta (4– 8 Hz) (REM) power percentage was assessed by calculating the power (FFT) per 10-s epoch and the average power per hour for every sleep epoch and finally were expressed hourly and in 72 h.

GSK1904529A, Chromeceptin administration and osmotic pump implantation.

The GSK1904529A and Chromeceptin was diluted in 5% DMSO, 20% PEG300 and 75% saline with final concentration of 0.5 mM and 1.5 mM respectively. The 5.5 mm stainless tube with 300 μm diameter was connected to a 100 μl osmotic pump through a rubber hose and then filled with DMSO/PEG300 (as control), GSK1904529A and Chromeceptin solution. mice were initially anesthetized in the isoflurane induction box (2.5% isoflurane) and placed in a stereotaxic head frame during surgery with 1.5% isoflurane. After exposure of brain bregma site, a stainless tube was then implanted with 3 degree and finally fixed above the SCN. Then all mice were recovered to cage with wheel after surgery. A heating pad was used to prevent hypothermia during surgery.

### Immunofluorescence and Nissl staining

Brains were collected, fixed in 4% paraformaldehyde overnight at 4 °C. Coronal sections (40 μm thick) were cut using vibratome (Leica VT1000s). Sections with middle SCN were used for IF. Free-floating sections were washed in PBS, incubated for 30 min in blocking buffer (1% BSA and 0.3% Triton X-100 in PBS) and incubated overnight at 4 °C with anti-KCC2 (19565-1-AP, Proteintech) primary antibody (abx129537, abbexa) diluted 1:1000 and 1:500 in the blocking buffer. Sections were then washed in PBS and incubated for 1 h in room temperature with donkey anti-rabbit Alexa-fluo (488) (Invitrogen) for KCC2 diluted 1:500 in blocking buffer. Sections were again washed, and incubated with DAPI (1:10,000 in PBS) for 5 min. After washed with PBS, sections were mounted on adhesion microscope slides (19017, Citotest) and air-dried in darkness. Slides were cover-slipped with 75% glycerinum and sealed with nail polish. Fluorescence-immunolabeled images were acquired using an Olympus fluorescence microscope.

For Nissl staining experiment, three 40 um coronal sections with anterior, medial and posterior part of DMH were incubated in cresyl violet in 37℃ for 10 min, then washed with H_2_O twice, 95% EtOH for 5S, 100% EtOH for 3 min, 50% EtOH plus 50% xylene for 5 min, xylene for 5 min, covered with resin. The images were acquired using an Olympus microscope.

### RNA-seq and differently expressed genes analyses

The SCN samples were collected with a 1.5mm tissue punch (Integra Miltex), and the total RNA was extracted with Trizol Reagent (Thermo). For RNA-seq, the quality of the total RNA was determined using an Agilent 2200 TapeStation, and RNA-seq was performed on an illumina NovaSeq 6000 platform with PE 150-bp reads at the Genewiz, Suzhou, China. Paired-end clean reads were aligned to the NCBI reference genome mm10 using Hisat2 (v2.0.5) and assembled using Stringtie (v1.3.3). Cuffdiff v1.3.0 was used to calculate FPKMs for coding genes in each sample.

To identify genes with a daily rhythmic expression from the gene expression data in different experimental groups, the Jonckheere-Terpstra-Kendall (JTK) algorithm was used. A permutation-based P-value (ADJ.P) of less than or equal to 0.05 was considered significant for all array sets. The optimal phase (LAG), amplitude (AMP), and period (PER) estimates for each transcript of oscillating genes were extracted from the JTK algorithm. Cuffdiff v2.2.1 provides statistical routines for determining differential expression in digital transcript or gene expression datasets using a negative binomial distribution model. Genes with corrected p-value less than 0.05 and the fold change greater than or equal to 1.5 were assigned as significantly differentially expressed genes (DEGs). Thus, rhythmic change and DEGs are defined as differently expressed genes.

### PCA and Gene Ontology analysis

RNA-seq raw data were first subjected to Bartlett’s Test of Sphericity to validate its adequacy for PCA (p < 0.0001), after which correlation-based PCA was implemented in R using factoextra and FactoMineR packages. Broken Stick model was used to determine the number of retainable PCs and we retained the first two PCs, which collectively explained 73.1% of the variance in different groups. Specifically, we aimed to identify the most relevant pathways to dawn behavior, the unsuitable pathways or enrichment annotations such as colorectal cancer, HIV infection, tuberculosis were removed from the datasets. Gene Ontology and pathway analysis were performed by DAVID (https://david.ncifcrf.gov/home.jsp) to significantly enriched GO terms and biological process at p values < 0.01 (Bonferroni) compared with the whole-transcriptome background.

### RT PCR

The SCN total RNA was extracted as mentioned before. For the quantitative real-time RT-PCR, the RNA concentration was determined by a NanoDrop (Thermo), and the cDNA was synthesized using a PrimeScript RT Reagent Kit (Takara). Quantitative real-time RT-PCR was performed using SYBR Premix Ex Taq with StepOne Plus (Applied Biosystems). The relative levels of *Kcc2* were normalized to *Gapdh*. The forward primer sequence for *Gapdh* was 5’-AGGTCGGTGTGAACGGATTTG-3’; the reverse primer sequence for *Gapdh* was 5’-TGTAGACCATGTAGTTGAGGTCA-3’. The forward primer sequence for endogenous *Kcc2* was 5’-GGGCAGAGAGTACGATGGC −3’; the reverse primer sequence for *Kcc2* was 5’-TGGGGTAGGTTGGTGTAGTTG −3’; the forward primer sequence for endogenous *Igf2* was 5’-GTGCTGCATCGCTGCTTAC −3’; the reverse primer sequence for *Igf2* was 5’-ACGTCCCTCTCGGACTTGG −3’.

### Statistics

The statistical analyses of food intake, body weight, locomotion range and slope were performed using t-tests using GraphPad Prism (GraphPad Software Inc., San Diego, CA, USA). A p < 0.05 was considered statistically significant (* p < 0.05, ** p < 0.01, and *** p < 0.001).

## Acknowledgements

We thank members of Cam-Su GRC for their assistance in the animal facility and Xu’s laboratory for discussion, and Hongbo Jia for his critical reading. This work was supported by grants from the National Science Foundation of China (31630091) to YX, the National Key R&D program of China (2018YFA0801100) to YX, and Science and Technology Project of Jiangsu Province (BZ2020067) to Y.X. We also thank the Priority Academic Program Development of the Jiangsu Higher Education Institutes (PAPD) and National Center for International Research (2017B01012). Vidal-Puig is supported by MRC MDU grant.

## Authors’ contributions

Q.Z. and Y.X conceived and designed the experiments. Q.Z conducted time-restricted feeding, all behavioral analysis, AAV stereotactic injection, fiber photometry recordings, Ca^2+^ signal data analysis, sleep recording and analysis, chemogenetic manipulation, SCN *Kcc2* knockdown and IGF2 intervention experiments. Q.Z. Y.G. Y.X. worked RNA-seq data analysis. Y.Z assisted with *Kcc2* knockdown experiment. Y.Y. assisted with tissue sampling. J.Y. and L.Y. provided helps in Ca^2+^ signal data analysis. H.Q. and X.C provided great helps in set up of fiber photometry system. Q.Z. A.V. P and Y.X. wrote the manuscript.

---

**Fig. S1.**
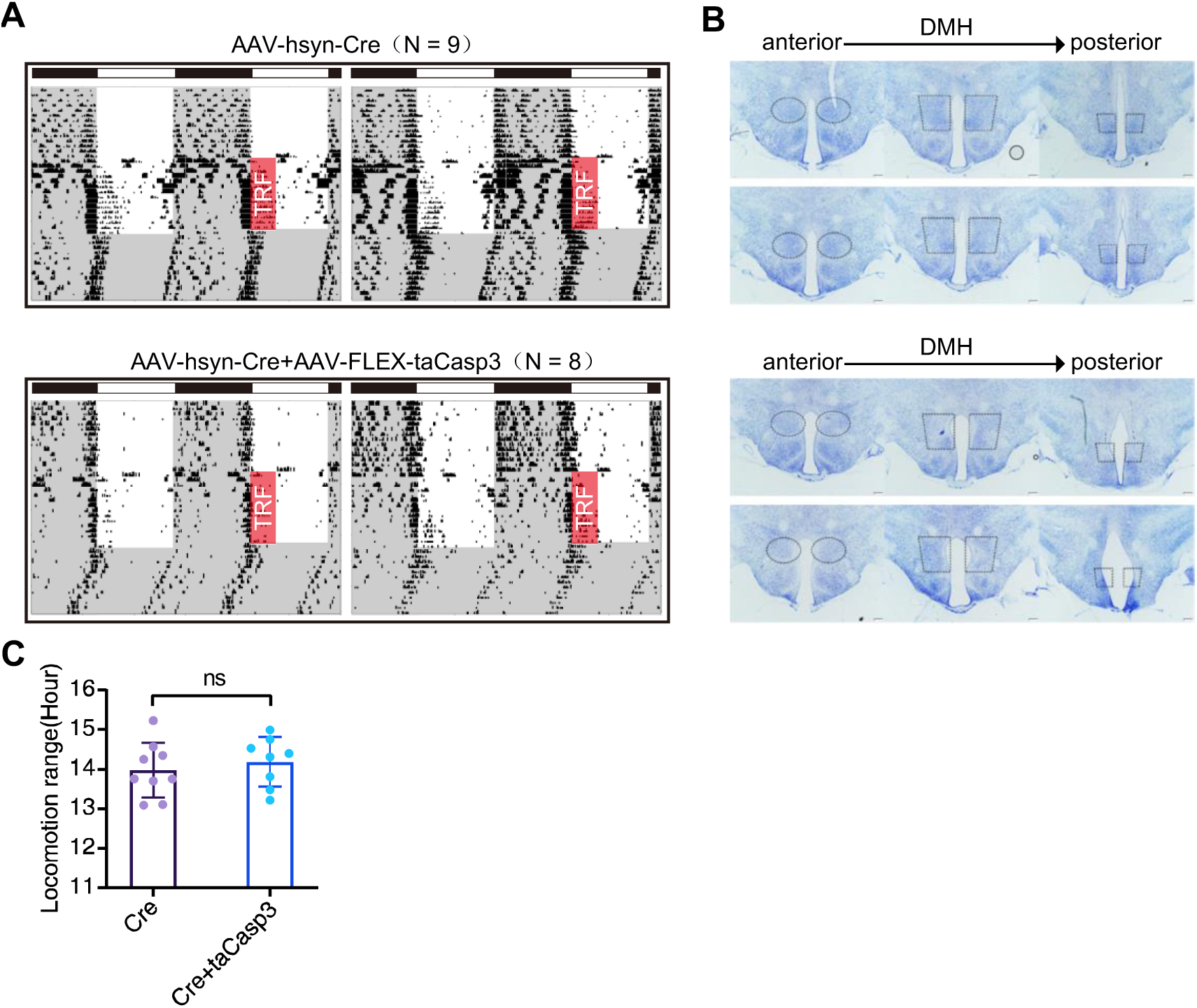
Mice with disruption of the DMH is unable to abolish the aftereffect of locomotor plasticity by ZT0-4 TRF. (**A**) The actograms of locomotors in the DMH lesioned mice with ZT0-4 TRF. (**B**) The lesioned areas in taCasp3-lesioned mice were confirmed by Nissl staining from anterior to posterior area of DMH. (Scale bar: 200 μm). (**C**) Statistics of the locomotion range from different groups. ns: not significant difference, Error bars: ± SD.

**Fig. S2.**
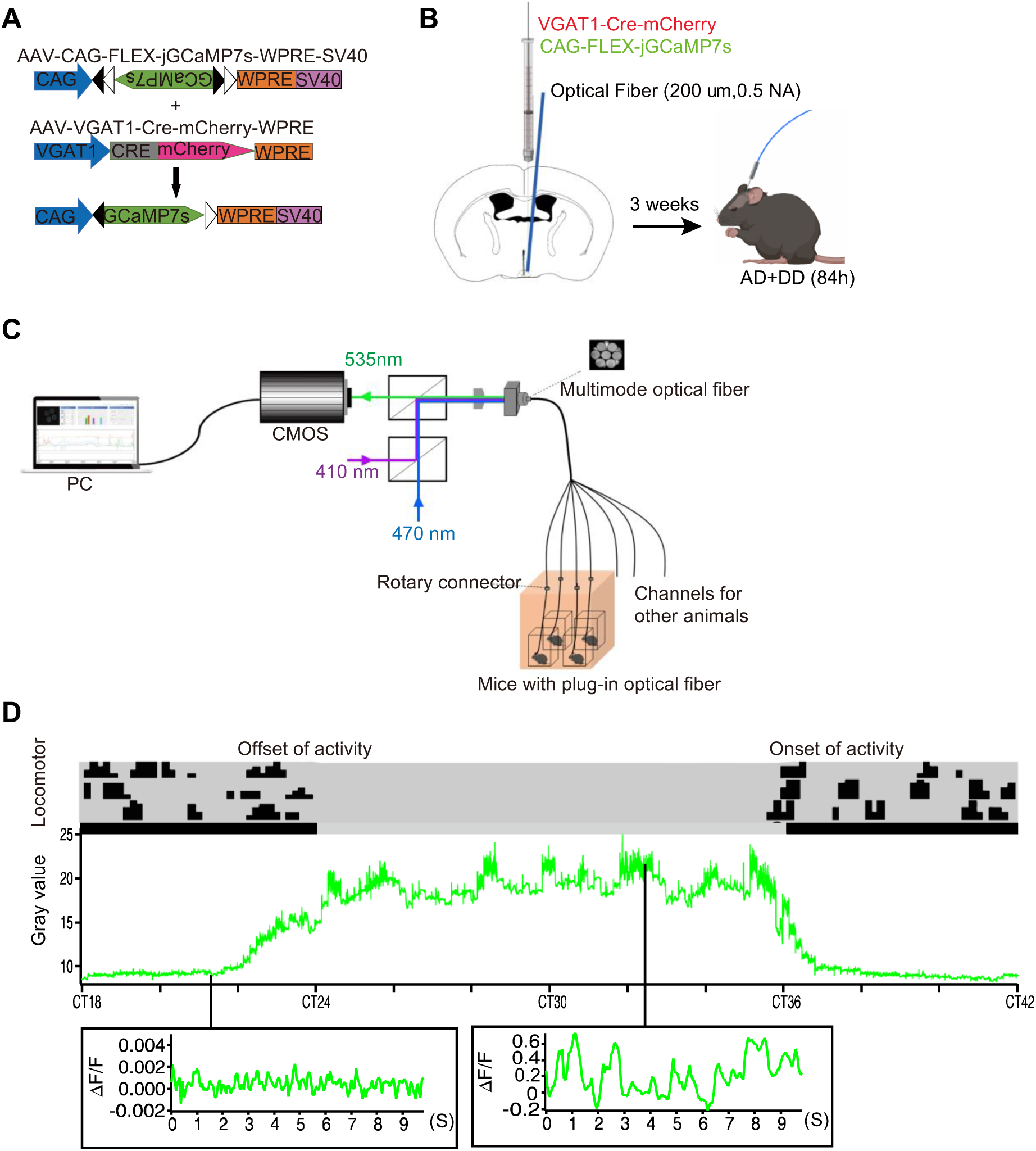
Established fiber photometry system to record neuronal activities in the SCN of freely-moving mice. (**A**) The skeleton of AAV-VGAT1-Cre and Cre inducible AAV-FLEX-jGCamp7s. (**B**) A mixture of AAV (150 nl) was injected in the SCN of C57/BL mice unilaterally followed by implantation of optical fiber (200 um,0.5 NA). Mice were allowed to recover for 3 weeks and then record Ca^2+^ signals. (**C**) The modules of fiber photometry system. Including a computer (PC) controlled CMOS photodetector structures, lasers with wavelength of 470 nm and 410 nm multimode optical fiber (7 channels), rotary connector, a box with controlled light/dark environment. Mice with plug-in optical fiber were housed in the box. (**D**) Ca^2+^ signals (raw fluorescence intensity) were recorded from ZT12 to CT72 under DD. A representative screenshot of Ca^2+^ signals from CT18-CT42 (n = 4). A zoom in signals from subjective night, while another is from the subjective day. A representative actogram of wheel-running was aligned to the Ca^2+^ signals. The ascending slope corresponds to offset of activity, and declining slope to the onset of activity.

**Fig. S3.**
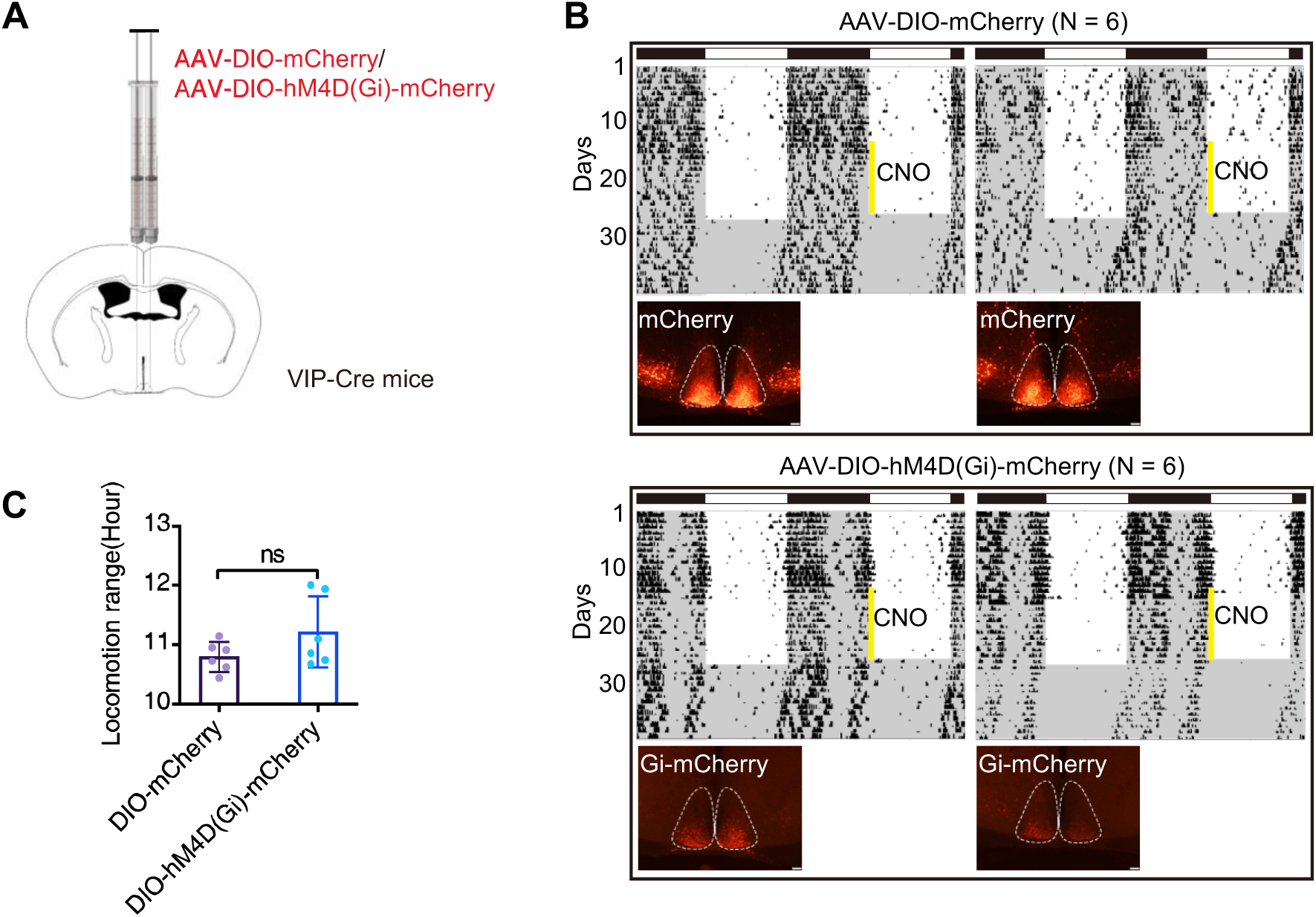
Inhibition of VIP neuron is unable to mimic the effect of TRF. (**A**) Cre dependent AAV of hM4D(Gi) and mCherry were injected in the SCN of VIP Cre mice. (**B**) The representative actogram of mice with intraperitoneal injected with CNO (3 mg/kg) for 13 days at ZT0 (yellow bar). (Scale bar: 100 μm). (**C**) Statistics of the locomotion range from different groups. ns: not significant difference, Error bars: ± SD.

**Fig. S4.**
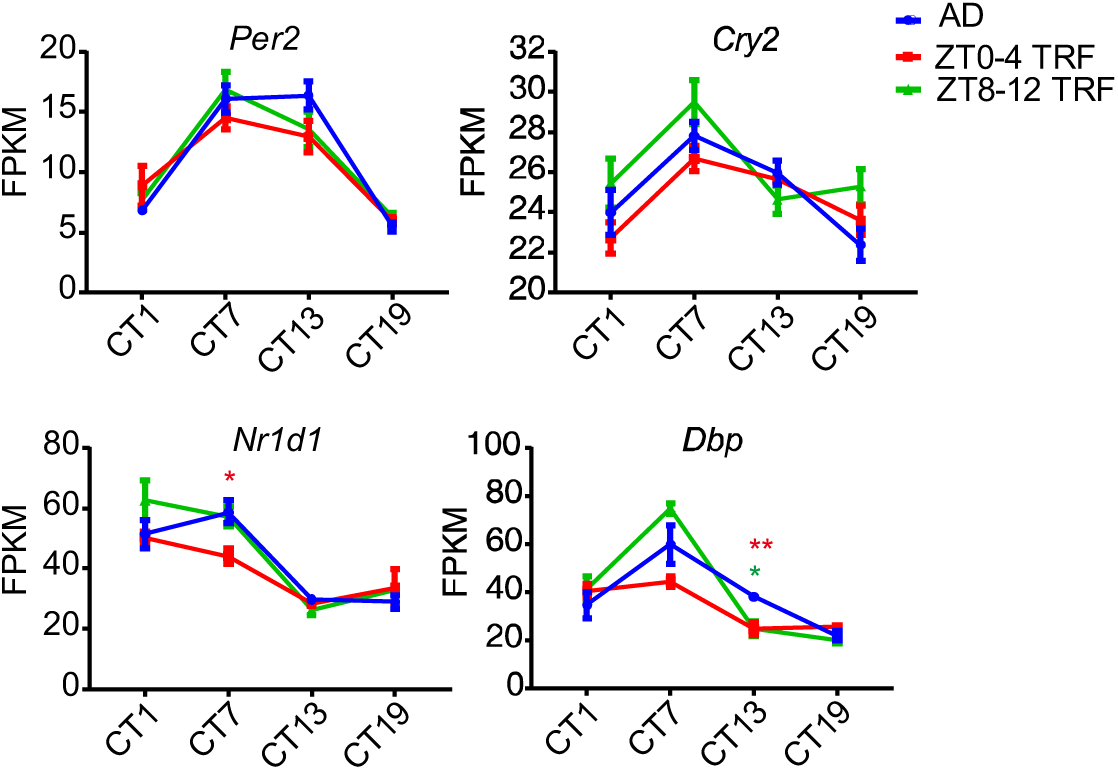
The FPKM of core clock gene. FPKM of clock genes (*Per2*, *Cry2*, *Nr1d1* and *Dbp*) in *ad libitum*, ZT0-4 TRF, and ZT8-12 TRF entrainment groups. Values represent the average ± SD. *: p < 0.05, **: p < 0.01. Red asterisk represents the difference between *ad libitum* and ZT0-4 TRF group, blue asterisk represents the difference between *ad libitum* and ZT8-12 TRF group.

## References

Abe, H., Honma, S., & Honma, K. (2007, Jan). Daily restricted feeding resets the circadian clock in the suprachiasmatic nucleus of CS mice. Am J Physiol Regul Integr Comp Physiol, 292(1), R607–615. https://doi.org/10.1152/ajpregu.00331.2006

Acosta-Galvan, G., Yi, C. X., van der Vliet, J., Jhamandas, J. H., Panula, P., Angeles-Castellanos, M., Del Carmen Basualdo, M., Escobar, C., & Buijs, R. M. (2011, Apr 5). Interaction between hypothalamic dorsomedial nucleus and the suprachiasmatic nucleus determines intensity of food anticipatory behavior. Proc Natl Acad Sci U S A, 108(14), 5813–5818. https://doi.org/10.1073/pnas.1015551108

Alvarez-Saavedra, M., Antoun, G., Yanagiya, A., Oliva-Hernandez, R., Cornejo-Palma, D., Perez-Iratxeta, C., Sonenberg, N., & Cheng, H. Y. (2011, Feb 15). miRNA-132 orchestrates chromatin remodeling and translational control of the circadian clock. Hum Mol Genet, 20(4), 731–751. https://doi.org/10.1093/hmg/ddq519

Ando, H., Ushijima, K., Shimba, S., & Fujimura, A. (2016, Feb). Daily Fasting Blood Glucose Rhythm in Male Mice: A Role of the Circadian Clock in the Liver. Endocrinology, 157(2), 463–469. https://doi.org/10.1210/en.2015-1376

Aton, S. J., Colwell, C. S., Harmar, A. J., Waschek, J., & Herzog, E. D. (2005, Apr). Vasoactive intestinal polypeptide mediates circadian rhythmicity and synchrony in mammalian clock neurons. Nat Neurosci, 8(4), 476–483. https://doi.org/10.1038/nn1419

Bass, J., & Takahashi, J. S. (2010, Dec 3). Circadian integration of metabolism and energetics. Science, 330(6009), 1349–1354. https://doi.org/10.1126/science.1195027

Challet, E. (2019, Jul). The circadian regulation of food intake. Nat Rev Endocrinol, 15(7), 393–405. https://doi.org/10.1038/s41574-019-0210-x

Cho, S., Muthukumar, A. K., Stork, T., Coutinho-Budd, J. C., & Freeman, M. R. (2018, Oct 30). Focal adhesion molecules regulate astrocyte morphology and glutamate transporters to suppress seizure-like behavior. Proc Natl Acad Sci U S A, 115(44), 11316–11321. https://doi.org/10.1073/pnas.1800830115

Choi, H. J., Lee, C. J., Schroeder, A., Kim, Y. S., Jung, S. H., Kim, J. S., Kim, D. Y., Son, E. J., Han, H. C., Hong, S. K., Colwell, C. S., & Kim, Y. I. (2008, May 21). Excitatory actions of GABA in the suprachiasmatic nucleus. J Neurosci, 28(21), 5450–5459. https://doi.org/10.1523/JNEUROSCI.5750-07.2008

Collins, B., Pierre-Ferrer, S., Muheim, C., Lukacsovich, D., Cai, Y., Spinnler, A., Herrera, C. G., Wen, S., Winterer, J., Belle, M. D. C., Piggins, H. D., Hastings, M., Loudon, A., Yan, J., Foldy, C., Adamantidis, A., & Brown, S. A. (2020, Aug 26). Circadian VIPergic Neurons of the Suprachiasmatic Nuclei Sculpt the Sleep-Wake Cycle. Neuron. https://doi.org/10.1016/j.neuron.2020.08.001

Crnko, S., Du Pre, B. C., Sluijter, J. P. G., & Van Laake, L. W. (2019, Jul). Circadian rhythms and the molecular clock in cardiovascular biology and disease. Nat Rev Cardiol, 16(7), 437–447. https://doi.org/10.1038/s41569-019-0167-4

Damiola, F., Le Minh, N., Preitner, N., Kornmann, B., Fleury-Olela, F., & Schibler, U. (2000, Dec 1). Restricted feeding uncouples circadian oscillators in peripheral tissues from the central pacemaker in the suprachiasmatic nucleus. Genes Dev, 14(23), 2950–2961. https://doi.org/10.1101/gad.183500

Escobar, C., Espitia-Bautista, E., Guzman-Ruiz, M. A., Guerrero-Vargas, N. N., Hernandez-Navarrete, M. A., Angeles-Castellanos, M., Morales-Perez, B., & Buijs, R. M. (2020, Apr 10). Chocolate for breakfast prevents circadian desynchrony in experimental models of jet-lag and shift-work. Sci Rep, 10(1), 6243. https://doi.org/10.1038/s41598-020-63227-w

Farajnia, S., van Westering, T. L., Meijer, J. H., & Michel, S. (2014, Jul 1). Seasonal induction of GABAergic excitation in the central mammalian clock. Proc Natl Acad Sci U S A, 111(26), 9627–9632. https://doi.org/10.1073/pnas.1319820111

Fernandez, A. M., & Torres-Aleman, I. (2012, Mar 20). The many faces of insulin-like peptide signalling in the brain. Nat Rev Neurosci, 13(4), 225–239. https://doi.org/10.1038/nrn3209

Fonken, L. K., & Nelson, R. J. (2014, Aug). The effects of light at night on circadian clocks and metabolism. Endocr Rev, 35(4), 648–670. https://doi.org/10.1210/er.2013-1051

Fuller, P. M., Lu, J., & Saper, C. B. (2008, May 23). Differential rescue of light- and food-entrainable circadian rhythms. Science, 320(5879), 1074–1077. https://doi.org/10.1126/science.1153277

Fuller, P. M., Lu, J., & Saper, C. B. (2009, Jul 22). Standards of evidence in chronobiology: A response. J Circadian Rhythms, 7, 9. https://doi.org/10.1186/1740-3391-7-9

Guerrero-Vargas, N. N., Espitia-Bautista, E., Buijs, R. M., & Escobar, C. (2018, Aug). Shift-work: is time of eating determining metabolic health? Evidence from animal models. Proc Nutr Soc, 77(3), 199–215. https://doi.org/10.1017/S0029665117004128

Hastings, M. H., Maywood, E. S., & Brancaccio, M. (2018, Aug). Generation of circadian rhythms in the suprachiasmatic nucleus. Nat Rev Neurosci, 19(8), 453–469. https://doi.org/10.1038/s41583-018-0026-z

Hastings, M. H., Maywood, E. S., & Brancaccio, M. (2019, Mar 11). The Mammalian Circadian Timing System and the Suprachiasmatic Nucleus as Its Pacemaker. Biology (Basel), 8(1). https://doi.org/10.3390/biology8010013

Herzog, E. D., Hermanstyne, T., Smyllie, N. J., & Hastings, M. H. (2017, Jan 3). Regulating the Suprachiasmatic Nucleus (SCN) Circadian Clockwork: Interplay between Cell-Autonomous and Circuit-Level Mechanisms. Cold Spring Harb Perspect Biol, 9(1). https://doi.org/10.1101/cshperspect.a027706

Houben, T., Deboer, T., van Oosterhout, F., & Meijer, J. H. (2009, Dec). Correlation with behavioral activity and rest implies circadian regulation by SCN neuronal activity levels. J Biol Rhythms, 24(6), 477–487. https://doi.org/10.1177/0748730409349895

Huang, W., Ramsey, K. M., Marcheva, B., & Bass, J. (2011, Jun). Circadian rhythms, sleep, and metabolism. J Clin Invest, 121(6), 2133–2141. https://doi.org/10.1172/JCI46043

Inagaki, N., Honma, S., Ono, D., Tanahashi, Y., & Honma, K. (2007, May 1). Separate oscillating cell groups in mouse suprachiasmatic nucleus couple photoperiodically to the onset and end of daily activity. Proc Natl Acad Sci U S A, 104(18), 7664–7669. https://doi.org/10.1073/pnas.0607713104

Jones, J. R., Simon, T., Lones, L., & Herzog, E. D. (2018, Sep 12). SCN VIP Neurons Are Essential for Normal Light-Mediated Resetting of the Circadian System. J Neurosci, 38(37), 7986–7995. https://doi.org/10.1523/JNEUROSCI.1322-18.2018

Kon, N., Yoshikawa, T., Honma, S., Yamagata, Y., Yoshitane, H., Shimizu, K., Sugiyama, Y., Hara, C., Kameshita, I., Honma, K., & Fukada, Y. (2014, May 15). CaMKII is essential for the cellular clock and coupling between morning and evening behavioral rhythms. Genes Dev, 28(10), 1101–1110. https://doi.org/10.1101/gad.237511.114

Kuhlman, S. J. (2007). Biological Rhythms Workshop IB: neurophysiology of SCN pacemaker function. Cold Spring Harb Symp Quant Biol, 72, 21–33. https://doi.org/10.1101/sqb.2007.72.061

Liu, Z., Huang, M., Wu, X., Shi, G., Xing, L., Dong, Z., Qu, Z., Yan, J., Yang, L., Panda, S., & Xu, Y. (2014, Jun 12). PER1 phosphorylation specifies feeding rhythm in mice. Cell Rep, 7(5), 1509–1520. https://doi.org/10.1016/j.celrep.2014.04.032

Mei, L., Fan, Y., Lv, X., Welsh, D. K., Zhan, C., & Zhang, E. E. (2018, Apr 17). Long-term in vivo recording of circadian rhythms in brains of freely moving mice. Proc Natl Acad Sci U S A, 115(16), 4276–4281. https://doi.org/10.1073/pnas.1717735115

Meredith, A. L., Wiler, S. W., Miller, B. H., Takahashi, J. S., Fodor, A. A., Ruby, N. F., & Aldrich, R. W. (2006, Aug). BK calcium-activated potassium channels regulate circadian behavioral rhythms and pacemaker output. Nat Neurosci, 9(8), 1041–1049. https://doi.org/10.1038/nn1740

Mistlberger, R. E., Buijs, R. M., Challet, E., Escobar, C., Landry, G. J., Kalsbeek, A., Pevet, P., & Shibata, S. (2009, Aug 10). Food anticipation in Bmal1-/- and AAV-Bmal1 rescued mice: a reply to Fuller et al. J Circadian Rhythms, 7, 11. https://doi.org/10.1186/1740-3391-7-11

Montgomery, J. R., Whitt, J. P., Wright, B. N., Lai, M. H., & Meredith, A. L. (2013, Feb 15). Mis-expression of the BK K(+) channel disrupts suprachiasmatic nucleus circuit rhythmicity and alters clock-controlled behavior. Am J Physiol Cell Physiol, 304(4), C299–311. https://doi.org/10.1152/ajpcell.00302.2012

Myung, J., Hong, S., DeWoskin, D., De Schutter, E., Forger, D. B., & Takumi, T. (2015, Jul 21). GABA-mediated repulsive coupling between circadian clock neurons in the SCN encodes seasonal time. Proc Natl Acad Sci U S A, 112(29), E3920–3929. https://doi.org/10.1073/pnas.1421200112

Nikhil, K. L., Korge, S., & Kramer, A. (2020, Aug). Heritable gene expression variability and stochasticity govern clonal heterogeneity in circadian period. PLoS Biol, 18(8), e3000792. https://doi.org/10.1371/journal.pbio.3000792

O’Neill, J. S., Maywood, E. S., Chesham, J. E., Takahashi, J. S., & Hastings, M. H. (2008, May 16). cAMP-dependent signaling as a core component of the mammalian circadian pacemaker. Science, 320(5878), 949–953. https://doi.org/10.1126/science.1152506

Olde Engberink, A. H. O., Huisman, J., Michel, S., & Meijer, J. H. (2020, Aug 31). Brief light exposure at dawn and dusk can encode day-length in the neuronal network of the mammalian circadian pacemaker. FASEB J. https://doi.org/10.1096/fj.202001133RR

Olde Engberink, A. H. O., Meijer, J. H., & Michel, S. (2018, Aug). Chloride cotransporter KCC2 is essential for GABAergic inhibition in the SCN. Neuropharmacology, 138, 80–86. https://doi.org/10.1016/j.neuropharm.2018.05.023

Oster, H., Damerow, S., Kiessling, S., Jakubcakova, V., Abraham, D., Tian, J., Hoffmann, M. W., & Eichele, G. (2006, Aug). The circadian rhythm of glucocorticoids is regulated by a gating mechanism residing in the adrenal cortical clock. Cell Metab, 4(2), 163–173. https://doi.org/10.1016/j.cmet.2006.07.002

Porcellati, F., Lucidi, P., Bolli, G. B., & Fanelli, C. G. (2013, Dec). Thirty years of research on the dawn phenomenon: lessons to optimize blood glucose control in diabetes. Diabetes Care, 36(12), 3860–3862. https://doi.org/10.2337/dc13-2088

Rohr, K. E., Pancholi, H., Haider, S., Karow, C., Modert, D., Raddatz, N. J., & Evans, J. (2019, Nov 20). Seasonal plasticity in GABAA signaling is necessary for restoring phase synchrony in the master circadian clock network. Elife, 8. https://doi.org/10.7554/eLife.49578

Saper, C. B., Lu, J., Chou, T. C., & Gooley, J. (2005, Mar). The hypothalamic integrator for circadian rhythms. Trends Neurosci, 28(3), 152–157. https://doi.org/10.1016/j.tins.2004.12.009

Schaap, J., Albus, H., VanderLeest, H. T., Eilers, P. H., Detari, L., & Meijer, J. H. (2003, Dec 23). Heterogeneity of rhythmic suprachiasmatic nucleus neurons: Implications for circadian waveform and photoperiodic encoding. Proc Natl Acad Sci U S A, 100(26), 15994–15999. https://doi.org/10.1073/pnas.2436298100

Shan, Y., Abel, J. H., Li, Y., Izumo, M., Cox, K. H., Jeong, B., Yoo, S. H., Olson, D. P., Doyle, F. J., 3rd, & Takahashi, J. S. (2020, Aug 4). Dual-Color Single-Cell Imaging of the Suprachiasmatic Nucleus Reveals a Circadian Role in Network Synchrony. Neuron. https://doi.org/10.1016/j.neuron.2020.07.012

Takahashi, J. S. (2017, Mar). Transcriptional architecture of the mammalian circadian clock. Nat Rev Genet, 18(3), 164–179. https://doi.org/10.1038/nrg.2016.150

Todd, W. D., Venner, A., Anaclet, C., Broadhurst, R. Y., De Luca, R., Bandaru, S. S., Issokson, L., Hablitz, L. M., Cravetchi, O., Arrigoni, E., Campbell, J. N., Allen, C. N., Olson, D. P., & Fuller, P. M. (2020, Sep 2). Suprachiasmatic VIP neurons are required for normal circadian rhythmicity and comprised of molecularly distinct subpopulations. Nat Commun, 11(1), 4410. https://doi.org/10.1038/s41467-020-17197-2

VanderLeest, H. T., Houben, T., Michel, S., Deboer, T., Albus, H., Vansteensel, M. J., Block, F. D., & Meijer, J. H. (2007, Mar 6). Seasonal encoding by the circadian pacemaker of the SCN. Curr Biol, 17(5), 468–473. https://doi.org/10.1016/j.cub.2007.01.048

Wang, T. A., Yu, Y. V., Govindaiah, G., Ye, X., Artinian, L., Coleman, T. P., Sweedler, J. V., Cox, C. L., & Gillette, M. U. (2012, Aug 17). Circadian rhythm of redox state regulates excitability in suprachiasmatic nucleus neurons. Science, 337(6096), 839–842. https://doi.org/10.1126/science.1222826

Welsh, D. K., Takahashi, J. S., & Kay, S. A. (2010). Suprachiasmatic nucleus: cell autonomy and network properties. Annu Rev Physiol, 72, 551–577. https://doi.org/10.1146/annurev-physiol-021909-135919

Zarrinpar, A., Chaix, A., & Panda, S. (2016, Feb). Daily Eating Patterns and Their Impact on Health and Disease. Trends Endocrinol Metab, 27(2), 69–83. https://doi.org/10.1016/j.tem.2015.11.007

Zhang, Z., Zhai, Q., Gu, Y., Zhang, T., Huang, Z., Liu, Z., Liu, Y., & Xu, Y. (2020, Aug 15). Impaired function of the suprachiasmatic nucleus rescues the loss of body temperature homeostasis caused by time-restricted feeding. Sci Bull (Beijing), 65(15), 1268–1280. https://doi.org/10.1016/j.scib.2020.03.025

